# Combinatorial optimization of mRNA structure, stability, and translation for RNA-based therapeutics

**DOI:** 10.1101/2021.03.29.437587

**Authors:** Kathrin Leppek, Gun Woo Byeon, Wipapat Kladwang, Hannah K. Wayment-Steele, Craig H. Kerr, Adele F. Xu, Do Soon Kim, Ved V. Topkar, Christian Choe, Daphna Rothschild, Gerald C. Tiu, Roger Wellington-Oguri, Kotaro Fujii, Eesha Sharma, Andrew M. Watkins, John J. Nicol, Jonathan Romano, Bojan Tunguz, Eterna Participants, Maria Barna, Rhiju Das

## Abstract

Therapeutic mRNAs and vaccines are being developed for a broad range of human diseases, including COVID-19. However, their optimization is hindered by mRNA instability and inefficient protein expression. Here, we describe design principles that overcome these barriers. We develop a new RNA sequencing-based platform called PERSIST-seq to systematically delineate in-cell mRNA stability, ribosome load, as well as in-solution stability of a library of diverse mRNAs. We find that, surprisingly, in-cell stability is a greater driver of protein output than high ribosome load. We further introduce a method called In-line-seq, applied to thousands of diverse RNAs, that reveals sequence and structure-based rules for mitigating hydrolytic degradation. Our findings show that “superfolder” mRNAs can be designed to improve both stability and expression that are further enhanced through pseudouridine nucleoside modification. Together, our study demonstrates simultaneous improvement of mRNA stability and protein expression and provides a computational-experimental platform for the enhancement of mRNA medicines.

## INTRODUCTION

Messenger RNA (mRNA) therapeutics hold the potential to transform modern medicine by providing a gene therapy platform with the capacity for rapid development and wide-scale deployment. Compared to recombinant proteins, manufacturing of mRNA is faster, more cost-effective, and more flexible because mRNA can be easily produced by *in vitro* transcription. Over the past decade, technological discoveries in the areas of mRNA modifications and delivery systems have rapidly advanced basic and clinical research in mRNA vaccines^1–3^. However, technical obstacles facing mRNA therapeutics are also apparent. For example, mRNA vaccines still suffer from decreased efficacy due to poor RNA stability in solution and *in vivo* and to limited expression of the payload mRNA; these are all pivotal issues that need to be carefully optimized for preclinical and clinical applications^3–6^.

The development of mRNAs redesigned for increased stability and expression could maximize the impact of producing, delivering, and administering therapeutic mRNAs. However, achieving such designs is hindered by a poor understanding of how the sequence and structure of an mRNA influence its expression and stability, both in solution and in cells. For example, it has typically been assumed that mRNAs with more stable secondary structure might have increased in-solution stability but would have lower in-cell protein output due to the increased difficulty of the cellular translation machinery to process through RNA structure^7^, but this has not been tested and some recent results suggest that there might not be such a tradeoff^8,9^. Our poor understanding of these design rules is due in part to the historical difficulty of rapidly synthesizing full-length mRNAs with different untranslated regions (UTRs) and coding sequences (CDSs) which would enable high-throughput experimental approaches comparing their stability and expression.

Here, we overcome these technical hurdles and characterize hundreds of full-length reporter constructs that encode mRNA sequences with a wide variety of UTRs and CDSs. We present a massively parallel reporter assay termed <u>P</u>ooled <u>E</u>valuation of m<u>R</u>NA in-solution <u>S</u>tability, and <u>I</u>n-cell <u>S</u>tability and <u>T</u>ranslation RNA-seq (PERSIST-seq), which enables systematic determination of the effects of UTR, codon choice and RNA structure on mRNA translation rates in human cells and on mRNA stability, both in cells and in solution. This represents the first large-scale screen of mRNA redesigned across its entire length. We further leverage the unique ability of the Eterna^10^ community, an online citizen science platform that enables participants to collectively solve RNA design puzzles, to devise solutions with high diversity in sequence and predicted structure. We integrate our datasets to develop a model that accurately predicts protein output for a given mRNA based on its ribosome load and in-cell stability. This model enables identification of optimal mRNA designs without the need to individually test all mRNAs for protein output. With the further aim of understanding the impact of mRNA structure on in-solution stability, we developed a high-throughput method termed In-line-seq to measure RNA degradation patterns arising from intrinsic in-line hydrolysis. Nucleotide-resolution data on thousands of small, structured model RNAs from the Eterna platform revealed novel design rules for reducing solution hydrolysis and enabled development of a regression model termed DegScore that enables *in silico* RNA sequence optimization for enhanced in-solution stability. Finally, we compare the effect of nucleoside modifications on mRNA performance, including unexpected results of pseudouridine (Ψ) and its derivatives on in-solution stability. Ultimately, our findings culminate in the fully automated design of mRNAs containing 5’ and 3’ UTR elements and structure-optimized CDS regions that simultaneously confer high stability in solution and high protein expression in cells. Together, the combination of optimized UTRs for expression, DegScore-optimized CDS structure, and Ψ modification provides a general technology that can be applied to stabilize and increase protein expression of candidate mRNA therapeutics. We envision that these design rules and our combinatorial mRNA optimization platform will be widely applicable to rapidly engineer future mRNA therapeutics that simultaneously optimize stability and potency.

## RESULTS

### A combinatorial library for systematic discovery of mRNA design rules

In search of design rules for stable and high-expressing mRNAs, we aimed to characterize a large number of mRNA sequence designs with extensive variations in 5’ UTR, CDS, and 3’ UTR regions. We took advantage of recent accelerations in commercial gene synthesis and developed the massively parallel assay PERSIST-seq (**Fig. 1A**, **Fig. S1**). In this method, mRNA variants can be assayed in parallel for translation efficiency in cells, for stability in cells, and for stability in solution. Full-length *in vitro* transcription (IVT) DNA templates were obtained through commercial gene synthesis services (Twist, Genscript, Codex). Each template incorporated three additional features: (1) a shared T7 promoter sequence for performing IVT, (2) barcodes in the 3’ UTR to enable multiplexing via inexpensive, short-read sequencing, and (3) a constant region at the 3’ end that enabled pooled PCR and reverse transcription (RT) reactions (**Fig. S1**). This design allowed one-pot amplification and analysis of the library using common flanking sequences. The library was *in vitro* transcribed, modified (3’ polyA-tailing and 5’ m^7^G-capping), transfected into cells and quantified by barcode sequencing in a pool (**Fig. 1B**), enabling straightforward measurements of translation by polysome profiling or of mRNA degradation over time in cells or in solution.

**Figure 1.**
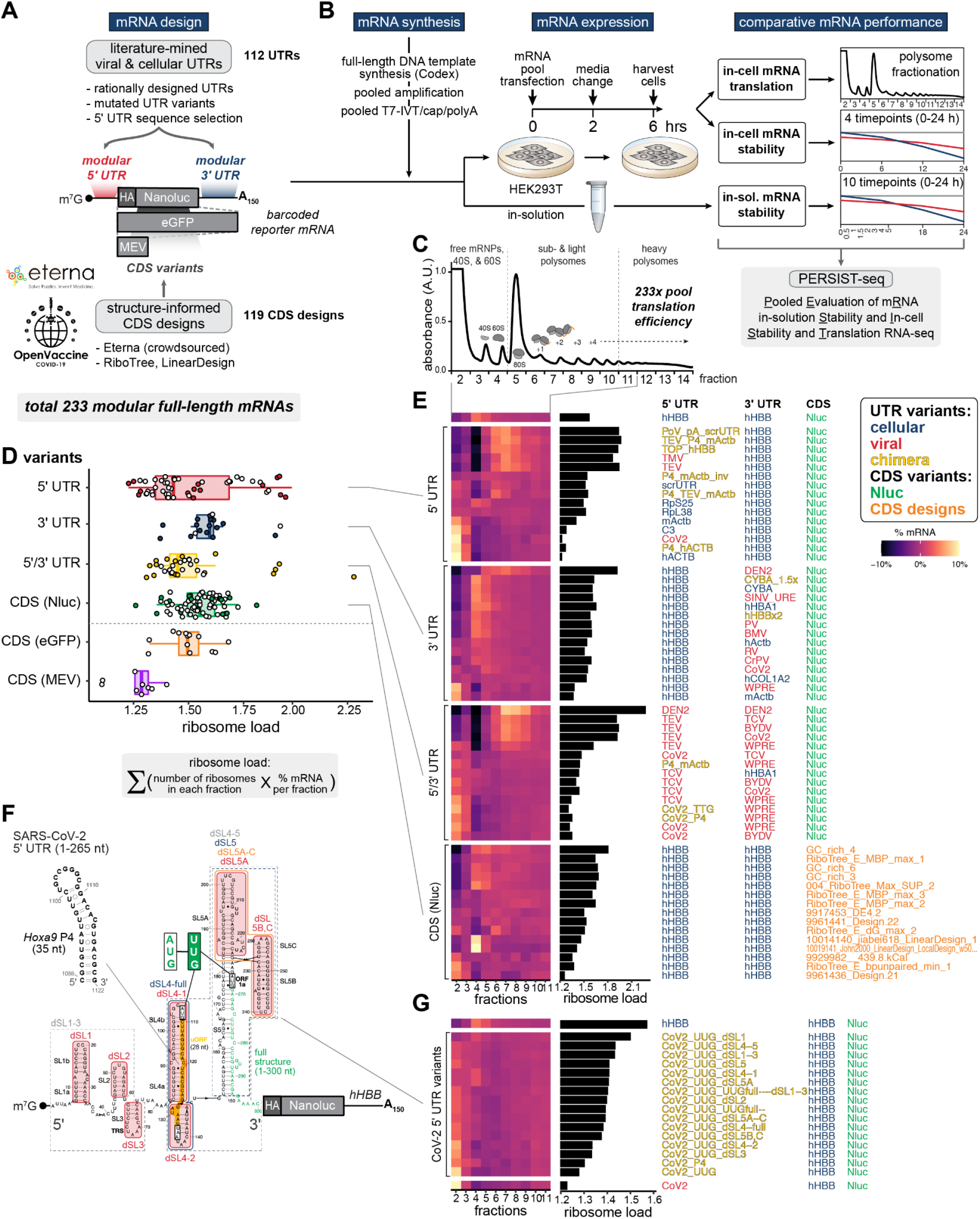
PERSIST-seq overview and illustrative ribosome load insights. (A) Overview of the mRNA optimization workflow. Literature mined and rationally designed 5’ and 3’ UTRs were combined with Eterna and algorithmically designed coding sequences. All sequences were then experimentally tested in parallel for in-solution and in-cell stability as well as ribosome load. The mRNA design included unique, 6-9 nt barcodes in the 3’ UTR for tag counting by short-read sequencing. (B) Experimental design for testing in-solution and in-cell stability and ribosome load in parallel. mRNAs were *in vitro* transcribed, 5’ capped, and polyadenylated in a pooled format before transfection into HEK293T cells or being subjected to in-solution degradation. Transfected cells were then harvested for sucrose gradient fractionation or in-cell degradation analysis. (C) Polysome trace from transfected HEK293T cells with 233-mRNA pool. (D) 5’ UTR variants display a higher variance in mean ribosome load per construct as determined from polysome sequencing. The formula for ribosome load is given. (E) Heatmaps from polysome profiles of mRNA designs selected from the top, middle, and bottom five mRNAs (by ribosome load) from each design category. (F) Secondary structure model of the SARS-CoV-2 5’ UTR. Introduced mutations and substitutions are highlighted. (G) Heatmaps of SARS-CoV-2 5’ UTR variants’ polysome profiles sorted by ribosome load.

Our mRNA library contained 233 different mRNA sequences in total (**Table S1**). 112 of these mRNAs contained varied 5’ and/or 3’ UTRs (**Fig. 1A**). While there have been many efforts to optimize protein expression by attaching UTRs found in highly translated or stable mRNAs^11–13^, previous work tested and characterized only a handful of candidates at a time, either with a focus on individual functional UTR elements^14^ or on screening randomized short UTRs^15–18^. We decided to harness full-length naturally occuring UTRs to optimize mRNA expression. Thus, we included a wide variety of 5’ and 3’ UTR sequences from cellular and viral genomes in our systematic analysis due to their potential to enhance or fine-tune mRNA translation or stability.

As examples of cellular sequences, we included short 5’ UTRs of cellular mRNAs corresponding to highly abundant proteins that have a high translation rate such as ribosomal proteins (RPs) (*RPS25*, *RPL31*, *RPL38*), or structural components such as *tubulin beta-2B chain* (*Tubb2b*) and *actin* (*ActB)*, as well as *human collagen, type I, alpha 2* (*hCOL1A2*)^19^. We further included regulatory 5’ UTR elements such as the 5’ terminal oligopyrimidine (TOP) motif from *RPL18*, which promotes translational activation downstream of mTOR^20,21^, as well as the *Hoxa9* P4 RNA stem-loop which functions as a translation enhancer^22^. 5’ UTRs previously identified in translation efficiency screens such as *complement factor 3* (*C3*), *cytochrome P450 2E1* (*CYP2E1*), and *Apolipoprotein A-II* (*APOA2*)^12^, as well as plant rubisco components (*RBCS3B*, *RBCS1A*)^23^, were also included. For 3’ UTR regions (total 22; size range 60-597 nt), we employed known stabilizing RNA structures such as the MALAT1 non-coding RNA 3’-stem-loop structure that resembles an expression and nuclear retention element (ENE) and engages a downstream A-rich tract in a triple helix structure^24,25^, as well as known expression enhancing 3’ UTRs such as those from *human hemoglobin subunit alpha 1* (*HBA1*)^26^ and *cytochrome B-245 alpha chain* (*CYBA*)^27,28^.

Viruses have evolved a suite of compact regulatory elements for hijacking the host translation machinery to effectively promote translation and stability of their own mRNAs. For example, internal ribosome entry sites (IRESs) can recruit ribosomes to initiate translation without the need for the full repertoire of eukaryotic initiation factors, whereas other elements in the 5’ and 3’ UTRs of viruses encode structures that facilitate long-range RNA-RNA interactions to enhance protein expression or mRNA stability. Therefore, several UTRs originating from viral genomes were included: the 5’ and 3’ UTR elements of the SARS-CoV-2 RNA genome^29,30^, along with variants described below; the dengue virus (DEN2) 5’ and 3’ UTRs, which are thought to enhance viral protein expression^31,32^; 5’ and 3’ UTR elements from various tombusviruses (e.g., turnip crinkle virus (TCV)) which encode 3’ cap-independent translational enhancer RNA structures that recruit the translational machinery^33,34^; tobacco mosaic virus^35^ (TMV) and tobacco etch virus^36^ (TEV) 5’ leader sequences; a poxvirus poly(A) leader sequence that is proposed to facilitate translation^37^; and the 3’ UTRs of Sindbis virus (SINV) and the rabies virus glycoprotein, which increase viral RNA stability through recruitment of host proteins^38–40^.

As our main reference, we chose the 5’ and 3’ UTRs from *human hemoglobin subunit beta* (*hHBB*), which is one of the most efficiently expressed mammalian mRNAs and is commonly used in investigations of mRNA translation and stability^41,42^. Non-*hHBB* UTRs are referred to here as “UTR variants.” To test these UTR variants, all reporter mRNAs encode the Nanoluc luciferase (Nluc) open reading frame (ORF) as its CDS region^43,44^. We decided to use Nluc because its short ORF of 621 nt allowed for synthesis and pooled amplification of full-length DNA templates with UTRs of up to 600 nt attached at each end. Employing Nluc further enabled precise quantitative readout for comparing translation efficiencies in follow-up experiments on individual mRNAs through ratiometric measurements including a firefly luciferase (Fluc) mRNA spike-in control in transfection experiments integrated with luciferase luminescence assays^22^. Additional mRNAs encoding for enhanced green fluorescent protein (eGFP) and a shorter candidate multi-epitope vaccine (MEV)^7^ were included as controls in some experiments, further discussed below.

To test the impact of CDS sequence and predicted CDS structure on mRNA stability and translation, we sought to maximize the diversity of CDS sequences and structures for model protein targets. Consequently, we asked participants from the Eterna massive open laboratory^10^ to design CDSs encoding a variety of model mRNAs, without specific optimization metrics, in a series of challenges (**Fig. 1A**). These puzzles included design challenges for eGFP, MEV, and Nluc (‘OpenVaccine: Design of eGFP and epitope mRNA molecules’ and ‘OpenVaccine: Lightning Round Design + Vote of Nanoluciferase’). Later rounds of design challenges included degradation-specific metrics (AUP and DegScore, discussed later) within the game interface to guide optimization. We also included CDSs using several algorithmic approaches. First, we included sequences designed using commercially available algorithms to optimize codon adaptation index (CAI)^45^. Second, we designed sequences using a “GC-rich” approach, in which each codon is stochastically sampled from codons highest in GC content, based on a strategy developed by CureVac researchers^9^. Third, we included CDSs designed using the LinearDesign algorithm^46^, which returns a deterministic minimal free energy solution that is weighted by codon optimality. Finally, we used the Ribotree Monte Carlo tree search method to optimize AUP for eGFP and to compare to eGFP designs developed by Moderna researchers^7,8^. These design methods yielded a total of 121 CDS variants in the library (**Fig. 1A**, **Table S1**). To ensure useful cross comparison, *hHBB* UTRs were used for each of these CDS variants. Together, all candidates from these diverse sources of UTR and CDS sequences were combined into a single library of 233 mRNA constructs.

### High dynamic range of translation driven by UTRs

To assess translation efficiencies relevant for potency of mRNA therapeutics, PERSIST-seq transfects mRNA pools into human cells (here HEK293T). Cell lysate then undergoes sucrose gradient fractionation, which separates mRNAs into actively translating and non-translating fractions that are analyzed by RT-PCR of barcode regions and Illumina sequencing. Actively translating mRNAs have a higher number of ribosomes associated with them and are found in polysomal fractions whereas non-translating or poorly translating mRNAs are present in the free mRNA fraction or are associated with 40S ribosomal subunits (**Fig. 1C**). After initial studies confirming differences in polysome loading of a highly translated endogenous mRNA, human *ActB*, with that of a transfected control mRNA that has scrambled short UTR sequences^43^ (**Fig. S1D**), we carried out PERSIST-seq to examine the polysome profiles of diverse constructs in the 233x-mRNA library. We observed a wide variation in mRNA distribution across the fractions (here expressed as ribosome load, defined as the weighted sum of mRNA proportions multiplied by the ribosome number in a fraction) (equation in **Fig. 1D**). The largest variation in ribosome load was observed for the 5’ UTR variants group (**Fig. 1D**). These data suggested strong potential for using different 5’ UTRs to tune translational efficiency of target mRNAs – more than for any other region (3’ UTR or CDS).

We categorized the origins of UTR sequences described above as “cellular”, “viral”, and “chimera” (modular UTR combinations). Overall, the mRNA designs with highest ribosome load were observed across 5’ UTRs of cellular as well as viral origins (**Fig. 1E**, **Table S1**). These 5’ UTRs included: mouse *COL1A2*, *Hoxa9 P4*, *Rpl18a TOP*, plant *RBCS1A*; the poxvirus poly(A) leader sequence fused to a scrambled 5’ UTR sequence and also the 5’ UTRs of plant viruses TEV and TMV. The dengue virus 5’ and 3’ UTRs both individually increase ribosome loading, and combining them into one mRNA resulted in an additive effect (**Fig. 1E**, **Table S1**). All of these sequences had a higher ribosome load (1.7-2.3) than the *hHBB* 5’ UTR (1.57), thereby identifying potential UTR design strategies to boost mRNA translational efficiency. Moreover, the chimeric fusion of the *hHBB* 5’ UTR with elements such as the TEV or 5’ TOP sequence in the same 5’ UTR increased polysome loading. Overall, our polysome screen successfully identified a wide range of 5’ UTR sequences that can be deployed to successfully promote translation within cells. Most surprising, in contrast to previous reports^47–49^, we found 5’ UTRs that are highly structured, such as the dengue virus (DEN2), can support efficient translation.

We next asked if the highly structured 5’ UTRs used in our screen could be augmented to further improve ribosome loading. To this end, we performed a detailed mutagenesis analysis of the structured 5’ UTR^30,50–52^ from SARS-CoV-2 genomic RNA^29,30^ as a paradigmatic example (**Fig. 1F**). We first observed that mutation of the uORF in the 5’ UTR (AUG mutated to UUG; CoV2-UUG) resulted in higher ribosome load than the wildtype 5’ UTR (**Fig. 1F-G**). Then, to systematically determine the impact of each stem-loop on ribosome load, we introduced partial and full truncations of stem-loops (SL) 1-5 (SL1-5) into CoV-2-UUG. (**Fig. 1F**, **Table S2**). Additionally, we included larger truncations of combined deletions of adjacent stem-loops and introduced the *Hoxa9* P4 stem-loop, a 35 nt element that recruits 40S ribosomal subunits to enhance translation^22^, in lieu of the uORF (**Fig. 1F-G**, **Table S2**). Intriguingly, polysome profiling of the 5’ UTR SL mutant constructs revealed a wide range of overall improved ribosome load (**Fig. 1G**). Deleting the full 5’ half (dSL1-3) or the 3’ half (dSL4-5) of the 5’ UTR overall increased ribosome load. In particular, deletion of SL1 from the SARS-CoV-2 5’ UTR increased ribosome load from 1.23 (CoV-2) to 1.5 (CoV-2-UUG-dSL-1) which almost reaches the level of ribosome load of *hHBB* (1.57) (**Fig. 1G**). These results on the SARS-CoV-2 5’ UTR indicate that ribosome load can be fine-tuned through the modulation of distinct elements in structured viral 5’ UTRs, and that these effects can be read out through PERSIST-seq.

Inspired by these PERSIST-seq results, we sought to further understand how sequence and structure variation of 5’ UTRs might modulate translation efficiency through an unbiased selection from a complex sequence library (**Fig. S2**). We selected for highly translating transcripts by transfecting an mRNA reporter library containing randomized 5’ UTR sequences and harvesting mRNAs associated with heavy polysomes (**Fig. S2A**). We further enriched these libraries for highly translating transcripts over five total rounds of selection and re-transfection of the heavily ribosome-loaded mRNAs from two independent starting pools (**Fig. S2A-C**). We then functionally assessed the protein output of the top 15 sequences by normalized read abundance (**Fig. S2D**). We found the most impactful effects of sequence k-mers at the 5’ and 3’ end of the randomized 5’ UTR stretch (**Fig. S2E**, **Table S3)**. These effects included significant depletion of 5’ UTRs that contain out-of-frame AUG start codons relative to the main ORF and enrichments of short stem-loop motifs promoting translation (**Fig. S2F**). Interestingly, one-by-one tests of the 5’ UTRs selected to have high ribosome load gave lower total protein output than the starting sequence (see **Supplemental Text**); this observation foreshadowed a tradeoff between ribosome load and mRNA stability that we dissect in greater detail below.

Beyond effects of varying sequence and structure of UTRs, we were interested in delineating how variations of the CDS might impact mRNA translation^3,8,45^. As noted above, we observed less variance in ribosome loading among CDS variants than for UTR variants (**Fig. 1D-E**). The only clear effects on ribosome load were that MEV-encoding CDS variants displayed less ribosome load than Nluc- or eGFP-encoding CDS variants, as expected from their shorter ORFs. For Nluc-encoding mRNAs, CDS variants were similar in ribosome load or showed a shift towards being less loaded with ribosomes. The ribosome load of CDS variants had no obvious correlation with the codon adaptation index (CAI), GC content, minimum free energy (MFE), addition of signal peptides, or nonsynonymous mutations (**Table S1**). We did note that mRNAs designed to have highly structured CDSs by the LinearDesign algorithm exhibited polysome fraction profiles dominated by 80S monosomes (**Fig. 1E**), further discussed below. Taken together with our results above, these PERSIST-seq measurements indicate that structured CDS regions and a wide variety of 5’ UTR elements from cellular and viral origins can sustain or even improve in-cell translation efficiency compared to reference mRNA sequences.

### In-cell mRNA stability is a major predictor of total protein output

The total protein output from an mRNA depends not only on its in-cell translation efficiency but also how long it remains intact once transfected into cells. To assess the in-cell mRNA stability of the library of constructs in a pooled fashion, PERSIST-seq quantifies the fractions of mRNAs remaining at multiple timepoints following transfection of the library into cells. To ensure recovery of intact full-length mRNAs rather than their degraded fragments, PERSIST-seq uses a two-step protocol, first generating amplicons covering the entire CDS regions of mRNAs through reverse transcription-PCR (RT-PCR) and then using primers flanking just the barcode region for a second PCR before short-read Illumina sequencing to count intact mRNAs per time point (**Fig S1**). Fits of an exponential decay function across the timepoints give an mRNA half-life (t_1/2_) for each library construct.

For our library of 233 mRNAs, PERSIST-seq gives a wide dynamic range of in-cell half-lives, ranging from less than 5 hours to over 15 hours (**Fig. 2A**). We had originally expected that the sub-library comprised of varying 3’ UTR sequences would give the most variation in in-cell stability, since those mRNAs included diverse *cis*-regulatory elements that are known to recruit cytoplasmic factors to aid or prevent mRNA decay^53–56^. However, surprisingly, we instead observed the widest in-cell stability variation in the CDS and 5’ UTR variant groups (**Fig. 2A**). We also noted that constructs with higher ribosome load values tended to be more unstable (**Fig. 2B**). More specifically, these unstable mRNAs include 5’ UTR or 5’/3’ UTR variants that were heavily polysome shifted (**Fig. 2B**, polysome/monosome ratio). Increased association to monosomes and thus a more moderate increase in overall ribosome load is positively associated with mRNA stability (**Fig. 2B**, monosomes/pre-polysome(pre-80S) ratios). Importantly, these findings identify an unexpected rule for the design of mRNA – excessively increasing translation efficiency may negatively impact mRNA stability. Stated differently, increased polysome load appears counterproductive to the goal of maximizing the total amount of protein expressed over time.

**Figure 2.**
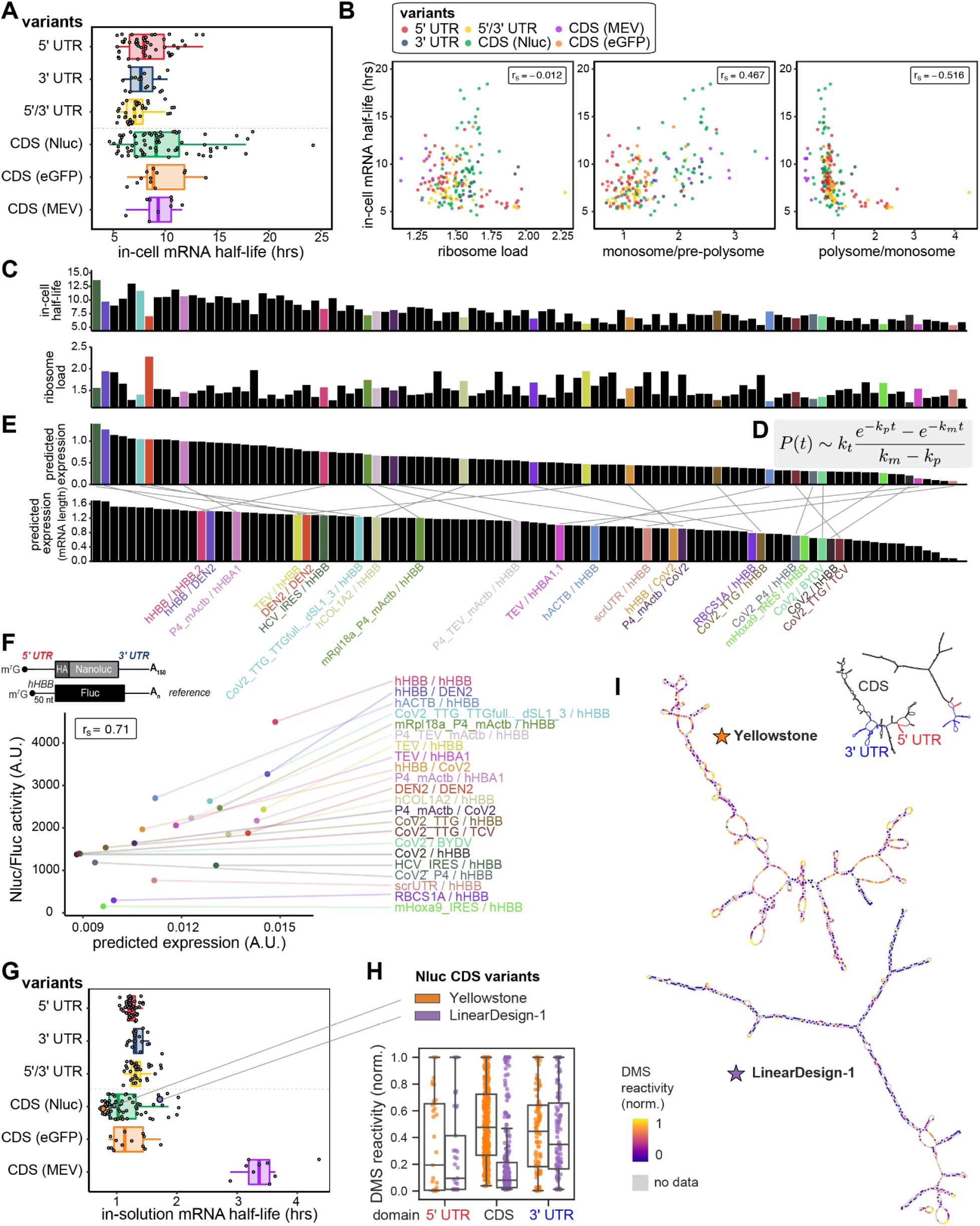
In-cell RNA stability drives downstream protein expression levels. (A) In-cell half-life of each mRNA design in HEK293T cells. (B) Higher polysome load correlates with decreased in-cell half-life. Correlation between in-cell half-life and mean ribosome load across the entire profile (left), monosome-to-free subunit ratio (center), or polysome-to-monosome ratio (right). (C) In-cell half-life and mean ribosome load for individual mRNA designs with varying UTRs. (D) Kinetic model for predicting protein expression from mRNA half-life and ribosome load. *P(t)* is protein quantity at time *t*; *m_0_* is the mass of mRNA present at *t=*0; *l* is mRNA length; *k_t_* is translation rate; and *k_m_* and *k_p_* are rates of mRNA and protein decay, respectively. (E) Protein expression predicted using the kinetic model in (D) on the basis of mRNA half-life and ribosome load. **Top**: predicted protein expression of each UTR variant; note closer similarity to in-cell half-life data than to ribosome load in (C). **Bottom**: predicted protein expression normalized by mRNA length (corresponding to transfecting equal masses of each mRNA). (F) Correlation of predicted protein expression and Nluc/Fluc activity in HEK293T cells. (G) In-solution half-life of various mRNA design variants. mRNA lifetimes are strongly dependent on mRNA length and designed structures, revealed by time courses of mRNA degradation under accelerated aging conditions (10 mM MgCl_2_, 50 mM Na-CHES, pH 10.0). (H) Nucleotide-resolution *in vitro* DMS mapping confirms large differences in structural accessibility between a highly structured JEV-HA-Nluc mRNA construct, “LinearDesign-1” and a highly unstructured construct “Yellowstone”. The 5’ and 3’ UTRs (*hHBB*) were kept constant between designs. (I) Nucleotide DMS accessibility mapped on to structures from DMS-directed structure prediction.

To explore the implications of this apparent tradeoff, we sought to integrate both translation (ribosome load) and in-cell stability (**Fig. 2C**) into a simple quantitative model to understand and predict their relative impact on integrated protein expression (**Fig. 2D-E**)^57^. To this end, we used differential equations to describe biochemical kinetics of mRNA translation and mRNA/protein decay assuming first-order rate of translation and exponential decay for mRNAs and for proteins. Below, *M(t)* and *P(t)* are moles of mRNA and protein at time *t*, respectively; *k_t_* is translation rate; and *k_m_* and *k_p_* are rates of mRNA and protein decay, respectively:

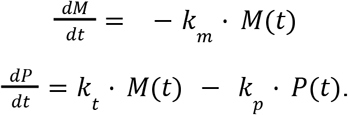

From these equations, the amount of protein at time *t* is proportional to the following (**Fig. 2D**):

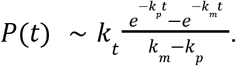

This expression enables the ranking of expected protein levels across the UTR variants in our mRNA library, as *k_t_* and *k_m_* are related to from PERSIST-seq measurements of ribosome load and in-cell half-lives, respectively, and the other parameters are known (**Fig. 2E**). Overall, predicted protein expression per mole of transfected mRNA is modeled to be mostly driven by the in-cell mRNA half-life (**Fig. 2C**, top panel), which more closely trends with predicted protein expression (**Fig. 2E**) than ribosome load does (**Fig. 2C**, lower panel). To test this model, we predicted and measured, using the luciferase readout, the protein levels observed at a given time (here 12 hours post-transfection) when an equal mass of mRNA is transfected for each sequence. Note that due to large length variations across tested UTR, the rank order of predicted protein output per equal mass of transfected mRNA is shifted (**Fig. 2E**, bottom panel). The predicted and measured luciferase activities showed a correlation of r = 0.71 (**Fig. 2F**), supporting the accuracy of our model. However, for short expression (early time points after mRNA transfection) or for “above-threshold” protein half-lives, translation can become the dominant predictor. Therefore, depending on the desired parameters (early burst of high protein expression or total protein output integrated over a longer time), the most optimal set of UTRs can vary. Taken together, our large-scale and unbiased pooled measurement of translation and mRNA stability provides a simple but powerful platform to test large numbers of different mRNA sequence designs for desired protein expression and has allowed us to importantly infer that in-cell mRNA stability is the dominant predictor of total protein output (**Fig. 2C,E**).

### mRNA length and structure drive in-solution mRNA stability

The degradation of RNA in solution is a major obstacle in the distribution of mRNA therapeutics to patients^6^. Thus, our final use of PERSIST-seq was to evaluate structure-based strategies for optimizing RNA designs to be more stable in solution. Just as with the in-cell stability measurements, we measured the fraction of mRNA CDS regions that remain intact after degradation, taking advantage of the same RT-PCR to select for intact mRNAs, followed by a PCR amplifying the short barcode regions in the 3’ UTR (**Fig. 1A**, **Fig. S1A, E**). To mimic the high effective pH and positively charged environment that can arise in lipid nanoparticles, protamine, and other formulations for mRNA therapeutics^6^, we used a high pH buffer containing Mg^2+^ to accelerate degradation (10 mM MgCl_2_, 50 mM Na-CHES, pH 10.0, 24 °C); conditions without Mg^2+^ or at lower pH lead to similar conclusions on relative stabilities across RNA variants (see below). The results for in-solution stability were strikingly different from the results for in-cell stability across the mRNA library. For example, in cells, modulation of UTR sequence produced large variation in in-cell stability (**Fig. 2A**), presumably through changes in recruitment of cellular machinery that affect mRNA decay. In contrast, in aqueous solution without such cellular machinery, changing the UTRs produced comparably little change in RNA stability to hydrolytic degradation of the CDS (**Fig. 2G**).

The largest changes in in-solution stability occurred across CDS variants. The strongest prediction of previous theoretical modeling^7^ was that length changes should drive the most variation in in-solution stability, and PERSIST-seq data across different different CDS types confirmed the effect of CDS length on RNA stability, (**Fig. 2G**). The shortest mRNAs in the pool, encoding a multi-epitope SARS-CoV-2 vaccine CDS (MEV), exhibited in-solution half-lives of 3.4±0.6 hours. The longest mRNAs, encoding eGFP, exhibited much shorter in-solution half-lives of 1.1±0.08 hours (**Fig. 2G**), as expected given the larger number of sites of potential hydrolysis. Indeed, the ratio of these half-lives, 3.0±0.7, matched within error the inverse ratio of the lengths of the mRNA regions captured by RT-PCR (958 nt/250 nt = 3.8), supporting theoretical predictions of length effects.

The next largest source of variation in in-solution stability was driven by differences in mRNA structure. Within mRNAs encoding for a single protein (Nluc, eGFP, or MEV), the variance in in-solution half-lives was greater than for the variance across UTR variants (**Fig. 2G**), and these values correlated well with different metrics for predicted structure (see below). The largest spread of in-solution half-lives (2.8-fold) occurred in the CDS variants for Nluc mRNAs. We chose two Eterna-submitted solutions amongst these mRNAs with short and long half-lives for follow up: ‘Yellowstone’, a design using codons that mimic the base frequencies (high A/C content) found in organisms found in the Yellowstone hot springs^58^; and ‘LinearDesign-1’, a design based on the LinearDesign mRNA structure optimization server^46^. Chemical structure mapping showed that the long-lived LinearDesign-1 was significantly more highly structured than Yellowstone, as assessed by dimethyl sulfate (DMS) and selective 2’-hydroxyl acylation with primer extension (SHAPE) reactivities^59,60^ (**Fig. 2H, Fig. S3**) and structure models guided by these data (**Fig. 2I**). Overall, the global assessment of in-solution RNA degradation using PERSIST-Seq reveals pronounced effects of RNA length and structure on RNA half-life in solution.

### Eterna-guided In-line-seq yields additional design principles and DegScore predictor

The mRNA designs above varied in-solution stability based mainly on computational predictions of mRNA structure and the assumption that nucleotides that are not base paired in structure would be uniformly prone to hydrolytic degradation^7^. We hypothesized that we might further improve in-solution stability through a deeper understanding of any specific sequence and structure features that lead to enhanced or suppressed hydrolysis in such unpaired regions. For example, base identities and local structural features such as the size and symmetry of apical loops and internal loops may play roles in determining in-solution mRNA degradation^61–63^.

To test such effects and to potentially discover new ones, we challenged Eterna participants to generate a large and diverse set of RNA molecules featuring designed secondary structure motifs in a special challenge (‘OpenVaccine: Roll-your-own-structure’, RYOS). Limiting the lengths of these molecules to 68 nucleotides and soliciting unique 3’ barcode hairpins enabled massively parallel synthesis and characterization of thousands of RNA molecules for their structure and degradation profiles (**Fig. 3A**). In particular, we obtained single-nucleotide-resolution measurements of 3030 RNA fragments using In-line-seq, a version of a low-bias ligation and reverse transcription protocol (MAP-seq)^64^ adapted here for in-line hydrolysis profiles^65^ (**Fig. 3B**). This is the first massively parallel methodology and large-scale data set applying in-line probing to RNA. For analysis of sequence and structure motifs, sequences were filtered for low experimental noise and to ensure that the structure predicted in ViennaRNA^66^ matched the structure inferred through SHAPE mapping data collected at the same time (see Methods); these filters resulted in 2165 sequences and corresponding secondary structures. We matched the accelerated degradation conditions used for PERSIST-seq in-solution stability measurements, but also verified that measurements without Mg^2+^, at lower pH, and higher temperature gave strongly correlated results (**Fig. S4A).** At a broad level, the data confirmed that RNA structure was a dominant predictor of in-line hydrolysis rate (compare, e.g., SHAPE to in-line data; **Fig. S4A**), but a closer look revealed additional sequence and structure dependent rules for in-line hydrolysis.

**Figure 3.**
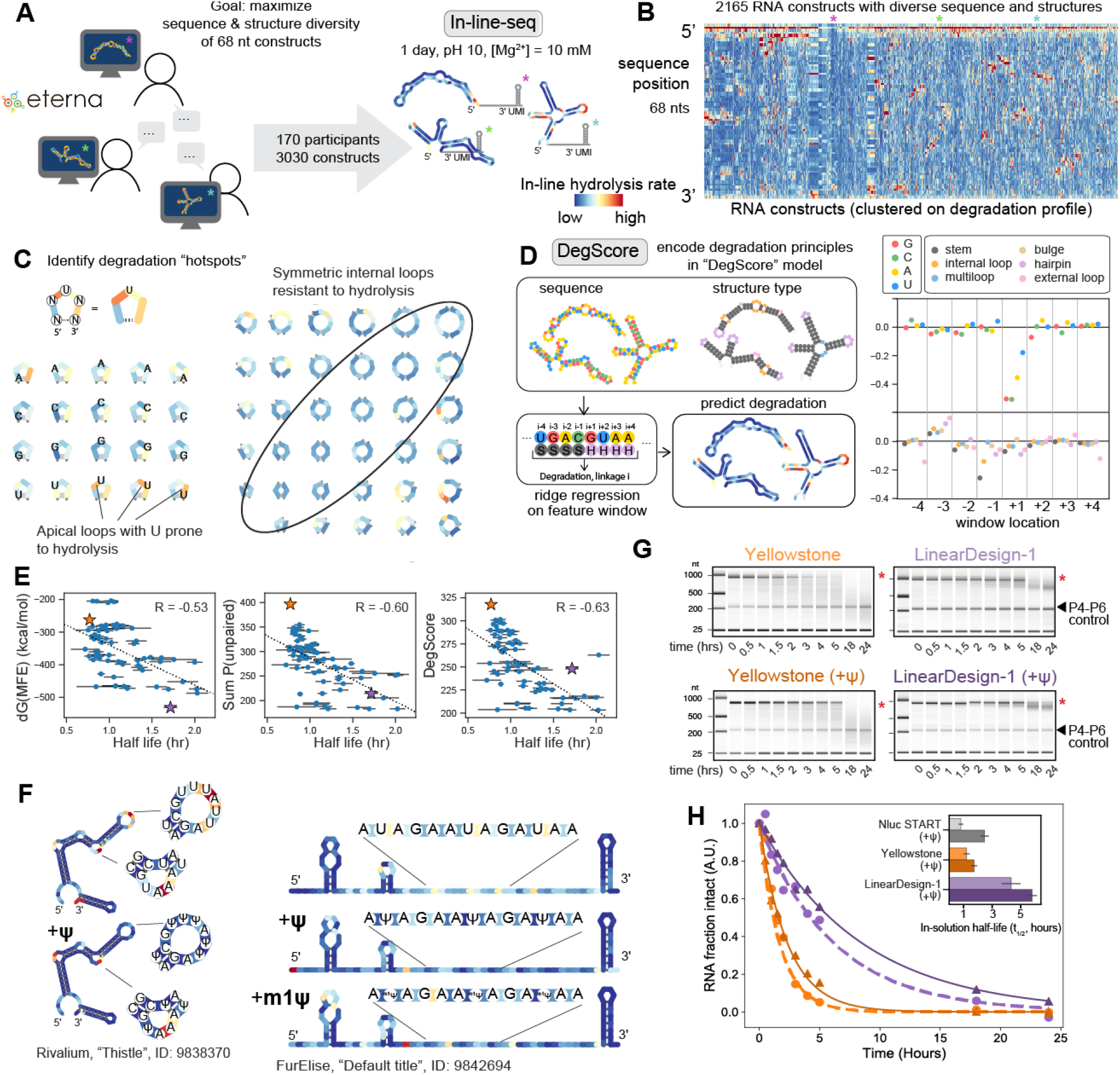
High-throughput in-line hydrolysis uncovers principles of in-solution RNA degradation. (A) Eterna participants were asked to design 68-nucleotide RNA fragments maximizing sequence and structure diversity. 3030 constructs were characterized and probed using high-throughput in-line degradation (In-line-seq). (B) Nucleotide-resolution degradation of 2165 68-nt RNA sequences (filtered for signal quality), probed by In-line-seq, sorted by hierarchical clustering on degradation profiles. (C) Sequences span a diverse set of secondary structure motifs, revealing patterns in degradation based on both sequence (i.e., linkages ending at 3’ uridine are particularly reactive) and structure (symmetric internal loops, circled, have suppressed hydrolytic degradation compared to asymmetric internal loops). (D) The ridge regression model “DegScore” was trained to predict per-nucleotide degradation from sequence and loop assignment information. Coefficients with the largest magnitude corresponded to sequence identity immediately after the link, with U being most disfavored. (E) DegScore showed improved predictive power on mRNAs over two other metrics previously posited to predict RNA stability. (F) Introduction of pseudouridine (Ψ) modifications stabilizes selected short RNAs at U nucleotides in both loop motifs and in fully unstructured RNAs. (G) Capillary electrophoresis characterization of fragmentation time courses of Nluc mRNA molecules designed with extensive structure (LinearDesign-1) and relatively less structure (Yellowstone), synthesized with standard nucleotides and with Ψ modifications. The full-length mRNA band is indicated with a red asterisk. The *Tetrahymena* ribozyme P4-P6 domain RNA was included after degradation as a control. (H) Exponential fits of capillary electrophoresis measurements of intact RNA over ten time points confirm >3x difference between in-solution lifetimes of LinearDesign-1 and Yellowstone Nluc mRNAs. Inset: Calculated half-lives. Error bars represent standard deviations from bootstrapped exponential fits (n = 1000).

When analyzed across known secondary structure motifs, the data revealed that the RNA sequence in a given structure can dramatically affect degradation of the structure motif. For example, in the case of the most-sampled secondary structure of triloops, the in-line hydrolysis rates varied by up to 100-fold depending on sequence (**Fig. S4B**). Furthermore, in many RNA loop types, it appeared that linkages that lead to a 3’ uridine were particularly prone to degradation (**Fig. 3C**, **Fig. S4C**), and this effect was reproduced in follow-up experiments using capillary electrophoresis readouts (**Fig. S4D**). Thus, independent of the nucleotide identity 5’ of the U, this bond is a hotspot for in-line nucleophilic attack^63,65^. In addition, we noted rules for hydrolytic degradation that depended on the type of RNA structural loop in which a nucleotide appears. A particularly salient characteristic was suppressed hydrolysis in symmetric internal loops compared to asymmetric internal loops (**Fig. 3C**). To distill these observations into a predictive model, we trained a windowed ridge regression model called ‘DegScore’ based on these In-line-seq data (**Fig. 3D**; see Methods) which quantitatively captured features like the increased hydrolysis rates at linkages leading to 3’ U (**Fig. 3D**). The DegScore regression coefficients with the largest magnitude corresponded to the identity of the nucleotide 3’ of a linkage (**Fig. 3D**). Of these coefficients, G and C were the most favorable (least hydrolysis) to have 3’ of the linkage, followed by A, and a 3’ U was most detrimental to degradation, matching our prior observations.

To test the accuracy of the DegScore metric derived from In-line-seq data, we made predictions of in-solution half-lives for the mRNAs measured in the PERSIST-seq experiments, which were carried out completely independently (**Fig. 1B**, **Fig. 2G**). For the Nluc CDS variants that showed the widest variance in in-solution half-lives, we observed a strong correlation of DegScore predictions to the in-solution half-lives (Pearson *R* = −0.63, *p*<0.0001). Strikingly, the accuracy of DegScore outperformed the accuracy of two other metrics that have been used in prior studies to parametrize RNA structure but do not take into sequence or structure-motif dependences of RNA hydrolysis: the free energy of the predicted minimum free energy secondary structure, the metric used in several design algorithms including LinearDesign (dG_MFE_; *R* = −0.53), and the predicted summed unpaired probability of the RNA structure ensemble^7^ (SUP; R= −0.60) (**Fig. 3E**). Beyond the Nluc CDS variants, we confirmed that DegScore gave the highest accuracy in predicting in-solution stability when evaluated over all the measured mRNAs, including low and high structure eGFP mRNAs from Moderna researchers, Eterna, and Ribotree (**Table S1** and **Fig. S5A,B**).

### Pseudouridine stabilizes RNA in solution

Given that the linkages with 3’ U were particularly sensitive to degradation, we hypothesized that the base chemistry of U may be directly linked to the degradative capacity of this nucleoside and sought to test whether chemical alternatives to U might alleviate degradation. In particular, we focused on Ψ and m^1^Ψ, since these substitutions for U have been widely adopted for mRNA therapeutics and vaccines due to improved in-cell translation and to better control of innate immune responses^2,3,8,67^ through avoidance of recognition by cellular toll-like receptors (TLR7 and TLR8)^68^, RIG‑I, and PKR^69–71^. While Ψ and derivatives have been reported to stabilize mRNAs against decay in cells^67^, the effects of these modifications on mRNA stability in solution have not been reported. We selected RNA sequences from the Eterna RYOS challenge (**Fig. 3A**) that were designed to contain U-rich loops or U-rich unstructured regions, resynthesized these RNAs one-by-one with standard nucleotides or with Ψ or m^1^Ψ substituted for U, and measured their in-line degradation over time via capillary electrophoresis. We observed that substitution of U with either Ψ or m^1^Ψ led to remarkable suppression of in-line hydrolysis at the substituted residues, presumably through changed nucleophilicity at the site of substitution (**Fig. 3F**). We also observed suppression of in-line hydrolysis at nucleotides 1 to 2 positions 5’ of the substitution (**Fig. 3F**), possibly due to locally enhanced base stacking^72^. Structure mapping data based on SHAPE and DMS profiling confirmed that Ψ and m^1^Ψ substitutions did not change the global structure of the RNAs; the suppression of in-line hydrolysis appears to be due to local chemical or structural effects.

As a further test of this unexpected stabilizing effect, we prepared six constructs from the 233x-mRNA library with and without Ψ (**Fig. 1**, **Fig. S5**), including the LinearDesign-1 and Yellowstone RNAs (**Fig. 2I**). In-solution degradation lifetimes were measured using capillary electrophoresis to evaluate the fraction of intact mRNA after different timepoints (**Fig. 3G-H**). Consistent with our in-line hydrolysis data on small RYOS RNAs, we observed a 1.2-2.7-fold stabilization for these longer Nluc-encoding mRNAs when U was uniformly substituted with Ψ (**Fig. S5C**). This finding indicated that beyond redesigning RNA sequences to take on stable structure, in-solution RNA stability can be significantly further improved by incorporating modified U nucleosides.

### One-by-one tests confirm additivity of UTR, CDS, and Ψ improvements

Thus far we found that mRNAs harboring highly structured 5’ and 3’ UTRs can support high levels of protein synthesis in cells. Moreover, using our RNA degradation predictor DegScore and Eterna-derived designs, we saw that the highly structured CDSs can strongly affect both the in-solution stability of mRNAs and their protein synthesis in cells (**Fig. 2**, **Fig. 3**). We next investigated if selected UTRs and CDSs might be combined to achieve stable and highly translated mRNAs that profit from additive effects of these individual improvements. In addition, we investigated the use of Ψ in place of uracils to more thoroughly determine its effects on stability and the overall translational output (**Fig. 4**).

**Figure 4.**
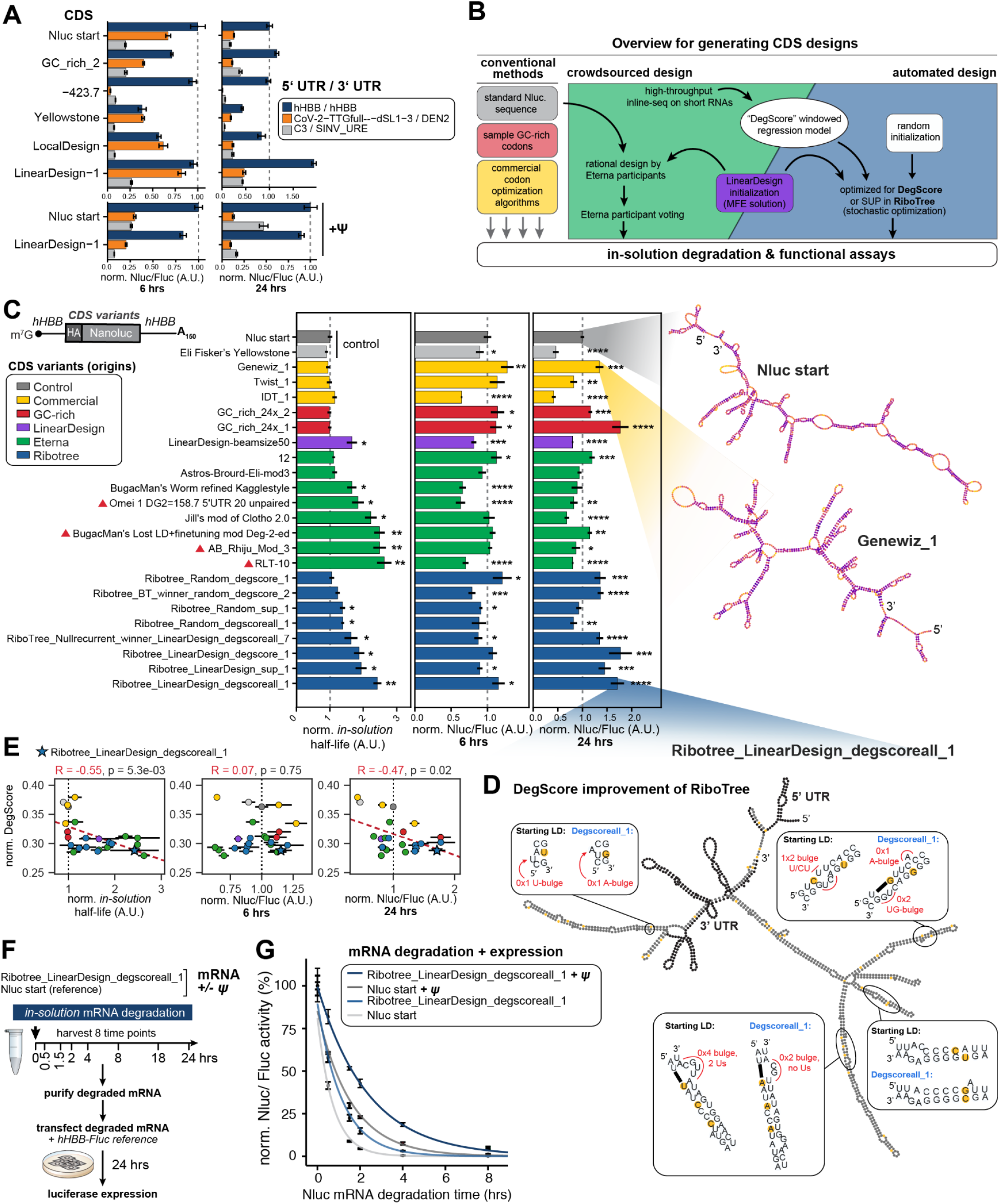
Integration of 5’/3’ UTRs, structure-optimized CDSs, and pseudouridine (Ψ) together enhance mRNA stability and translational output. (A) CDS and 5’/3’UTR combinations differentially impact protein synthesis. Six mRNA constructs were *in vitro* synthesized and luciferase activity was measured 6 or 24 hrs post-transfection. Inclusion of Ψ was tested on two selected constructs. (B) Workflow for different approaches to design the CDS variants tested in (C). (C) Variations in CDS design facilitates high in-solution stability and differential protein expression. *In vitro* transcribed mRNAs (24 in total) were subjected to *in-solution* degradation or transfected into HEK293T cells for 6 and 24 hrs. In-solution half-lives and luciferase activity are normalized to the Nluc START reference construct. Predicted secondary structures are shown for select constructs with colors indicating DegScore at each nucleotide. Designs derived from LinearDesign solutions are marked with a purple triangle. (D) Predicted secondary structure overview of Ribotree_LinearDesign_degscoreall_1. Zoomed boxes indicate sequence optimizations and subsequent structural changes made by DegScore to the reference LinearDesign construct. (E) Increased in-solution half-life and enhanced luciferase expression at 24 hrs, but not 6 hrs, correlate with DegScore. (F) Schematic for testing the synergy between RNA modifications and mRNA design rules on downstream stability and protein output. mRNAs were *in vitro* synthesized with or without Ψ and subjected to degradation conditions. Samples were collected overtime and the RNA was purified before being transfected into HEK293T cells. Luciferase activity was measured 24 hrs after transfection. (G) Luciferase activity of the reference Nluc sequence and DegScore-optimized CDS with or without Ψ after being subjected to in-solution degradation.

We first determined the combined contribution of different CDS designs and UTRs on mRNA stability and protein expression by choosing six CDS designs and three UTR combinations from our high-throughput screen (**Fig. 1A**, **Fig. 4A**). The selected CDS designs represented CDSs from diverse origins that support a range of observed in-solution half-lives (from 0.69- to 1.8-fold relative to our reference ‘Nluc start’ sequence; **Fig. 2G**). We combined these CDSs with different 5’ and 3’ UTRs in combination that were individually predicted and/or confirmed (**Fig. 2F**) to facilitate the highest protein expression in our library (**Fig. 2E-F**). The three UTR combinations comprised our standard *hHBB* 5’ and 3’ UTRs; a SARS-CoV-2 5’ UTR dSL-3 variant paired with the dengue virus 3’ UTR, predicted to have high translational efficiency but shorter in-cell half-life due to increased length; and C3 5’ UTR paired with a SINV U-rich element 3’ UTR predicted to have good translational efficiency while reducing the size of flanking sequence (**Fig. 2E-F**, **Table S1**). In terms of in-solution stability, we expected the mRNAs with the longer UTRs to have consistently reduced half-life to hydrolytic degradation across all 6 CDSs, and capillary electrophoresis experiments confirmed this expectation (**Fig. S6**).

To quantitatively evaluate in-cell protein expression, the 18 mRNAs were individually transfected and luciferase activity was measured after 6 hours (as an assessment of translation rate before significant mRNA decay) and 24 hours (as an assessment of total protein output after mRNA decay) (**Fig. 4A**). After 6 hours, two CDSs (LinearDesign-1 (**Fig. 3**) and GCrich_2) achieved similarly high protein levels compared to Nluc start, when combined with *hHBB* 5’ and 3’ UTRs. The LinearDesign-1 result was striking given its unusual monosome-concentrated polysome profile in PERSIST-seq (**Fig. 1E**) and the expectation that high mRNA structure should adversely impact the cellular translational apparatus. The result was nevertheless consistent with our model that enhanced in-cell half-life of a structured mRNA can compensate for low translation efficiency (**Fig. 2C-E**). Indeed, by 24 hours, the mRNA with the LinearDesign-1 CDS displayed an outstanding two-fold increase in luciferase yield compared to that of Nluc start (**Fig. 4A**). LinearDesign-1 CDS mRNAs also exhibited a particularly long in-solution half-life (**Fig. 3G-H**). Among the three UTR combinations and six CDS designs tested, most demonstrated lower overall luciferase activity than the reference start Nluc with *hHBB* UTRs (**Fig 4A**). One exception was the CoV-2-UUG-dSL-3/DEN2 UTR combination, chosen based on its high performance for ribosome load (**Fig. 2D-E**), which was able to support levels of protein synthesis at 6 hours nearly as high as the *hHBB* UTR for the LinearDesign-1 CDS; however this expression was lowered by 24 hours (**Fig 4A**). This finding is consistent with faster mRNA decay compared to *hHBB* UTRs and our results supporting that in-cell mRNA stability is a primary driver of protein output (**Fig. 2C, E**).

Given the improvements in in-cell stability we observed for pseudouridine (Ψ)-modified RNA fragments and Nluc mRNAs, we further tested the effect of pseudouridylation on in-cell stability and protein levels achieved from Nluc start vs. LinearDesign-1 CDS (**Fig. 4A**). As expected, preparation of mRNAs with Ψ substituted for U led to increased in-solution stability across both these CDSs, independent of UTR (**Fig. S6**). In terms of protein expression, we observed variable effects on overall luciferase activity in the different UTR combinations with fixed CDSs compared to unmodified mRNAs after both 6 and 24 hours of expression. Importantly, both Nluc start and LinearDesign-1 CDSs with *hHBB* UTRs maintained high protein expression at 6 and 24 hours (**Fig. 4A**) indicating that, despite Ψ modification and highly structured CDSs, translation is sustained. Overall, these results demonstrated the importance of mRNA stability to protein output, and suggested that the *hHBB* UTRs, highly structured CDSs, and use of Ψ would be the best choices for increasing in-solution stability and protein output moving forward.

### Integration of all design rules leads to mRNAs with both high in-solution stability and high protein output

For our final experiments, we sought to test whether further optimization, especially of the CDS, might allow both enhanced in-solution mRNA stability and in-cell protein expression. To this end, we collected a variety of CDS designs to compare to the Nluc start and Yellowstone mRNAs from above, including (1) the default output of available mRNA design algorithms, including those provided by Genewiz, Twist, and IDT websites and others that may enhance mRNA structure (LinearDesign^46^ and use of GC-rich codons^9^), (2) highly structured constructs that were rationally designed through an additional Eterna competition (‘OpenVaccine: Focus on the NanoLuciferase mRNA’), and (3) output of the automated mRNA structure design tool Ribotree, a stochastic optimization algorithm that can start from different seed sequences (random or LinearDesign) and optimize in-solution hydrolysis lifetimes as guided by different predictors (AUP, DegScore) (**Fig 4B**). From these approaches we generated 24 different CDS designs for Nluc mRNA and, based on our studies above, we added the *hHBB* 5’ and 3’ UTRs as constant regions to mediate high rates of translation initiation and used Ψ during synthesis due to its enhancement of in-solution mRNA stability (**Fig. 3F-G**).

Each mRNA design was subjected to accelerated degradation and capillary electrophoresis to measure the in-solution half-life of each individual CDS. Compared to the Nluc start reference sequence, designs originating from Eterna exhibited up to 2.6-fold higher in-solution half-life (see, e.g., RLT-10) while designs from commercial algorithms or optimized GC-content tended to exhibit similar in-solution half-lives (**Fig. 4C**). Designs from the LinearDesign server and modifications to these designs from both Eterna participants (e.g., AB_rhiju_mod3) and the RiboTree algorithm produced constructs with increased half-life up to approximately 2.5-fold over the Nluc reference sequence (designs marked with triangles in **Fig. 4C**). In parallel to measurements of in-solution half-life, individual mRNA designs were assayed for protein expression. Interestingly, despite overall longer in-solution half-lives, at 6 hours, both Eterna- and Ribotree-derived designs tended to have similar or lower luciferase activities than the reference Nluc sequence, whereas most vendor-derived and GC-rich designs had slightly higher activities (**Fig. 4C**). At 24 hours, the trend of lower luciferase activity continued for 6 of 8 Eterna-derived designs tested compared to the Nluc reference mRNA (**Fig 4C**, green). However, in contrast, 6 of 8 RiboTree-optimized mRNA designs demonstrated higher luciferase activities than the Nluc reference mRNA at 24 hours (**Fig. 4C**, blue). Remarkably, the sequence outputted by RiboTree starting from a LinearDesign CDS solution, with optimization guided by DegScore and taking into account the flanking *hHBB* UTR sequences, yielded an mRNA that was both highly stable in solution (t_1/2_ = 2.4, relative to Nluc start) and exhibited high levels of protein expression in cells (1.7-fold increase, relative to Nluc start) (Ribotree_LinearDesign_degscoreall_1; **Fig. 4C**). This simultaneous increase in in-solution stability with improved and sustained in-cell protein expression provided a strong demonstration of the impact of our design rules for mRNA.

To gain insight into what led to success for this RiboTree_LinearDesign_degscoreall_1 sequence, we examined what changes RiboTree made as it computationally minimized the DegScore metric from a starting LinearDesign server solution. Specific regions that RiboTree modified from the starting sequence are indicated in **Fig. 4D** (with further comparison in **Fig. S7**). These computational modifications are characterized by reducing the presence of Us in loops, and shifting local base-pairing to minimize the overall size of loops, even if such shifts result in additional smaller loops. These modifications are consistent with mitigating hydrolysis as modeled in the DegScore predictor **(Fig. 3D**). Taken together, these data suggest that by reducing the overall presence of Us in loops and reducing the number of hairpins to generate a “linear” highly double-stranded mRNA can lead to enhanced mRNA stability and protein expression.

Correlating data from these final 12 Nluc RNAs provided additional insight into biophysical and biophysical features impacting mRNA performance (**Fig. 4**, **Fig S8**). Most notably, DegScore correlated strongly with measured in-solution half-life (*R* = –0.55, *p*<0.001), and moderately with 24 hour protein expression (*R* = –0.47, *p* = 0.02), yet was uncorrelated with 6 hour protein expression (**Fig. 4E**). However, 6 hour protein expression correlated strongly with the predicted number of hairpins (*R* = 0.70, *p*<0.001) and the “Maximum Ladder Distance^73^,” or the maximum helix path length (*R* = −0.55, *p*<0.001) (**Fig. S8C**, **Table S5**). These observations suggest that resistance to RNA hydrolysis, as predicted by DegScore and quantified by in-solution half-life, is important for longer protein expression, but that other RNA sequence and structural features govern protein expression at shorter timescales. For instance, longer or more branched double-stranded RNA stems may potentially hinder the ability of the ribosome to unwind RNA secondary structure.

In all our tests above, we measured in-solution half-lives of mRNAs using structural readouts (reverse transcription-PCR; capillary electrophoresis), but sustaining functional output after degradation is of strongest interest for mRNA applications. As a final experiment, we therefore carried out an experimental stress test of in-solution stability with a functional readout based on transfection and protein production in cells. For this ‘end-to-end’ test of mRNA efficacy, we synthesized the original Nluc reference (Nluc start) mRNA alongside the optimized Ribotree_LinearDesign_degscoreall_1 mRNA both with and without Ψ. Each individual Nluc mRNA was then subjected to solution degradation, with 8 time points collected (**Fig. 4F**). As expected from the characterization above (**Fig. 4C**), the optimized Ribotree_LinearDesign_degscoreall_1 mRNA exhibited higher resistance to degradation in-solution than the Nluc Start mRNA and incorporation of Ψ into either mRNA further enhanced in-solution stability, this time with a readout based on mRNA function in cells (**Fig 4G**). The overall difference is striking: the majority of the mRNA stabilized with Ψ and a structure-optimized CDS remains functional after 2 hours of accelerated solution degradation, whereas our starting mRNA sequence gives negligible in-cell activity after the same degradation (**Fig 4G**). Taken together, our detailed studies on individual mRNAs (**Fig. 4**) indicate that combining UTRs that facilitate high levels of translation alongside structural optimization of CDSs via LinearDesign and DegScore-guided RiboTree design, we can enhance both in-solution stability and total in-cell protein output of mRNAs. Moreover, downstream luciferase expression can be further amplified by Ψ modification of mRNAs. These results validate and demonstrate a modular and flexible platform that is applicable to potentially any protein target of choice and can accelerate the design of overall improved mRNA therapeutics solutions.

## DISCUSSION

mRNA-based therapeutics are transforming the way in which human disease is treated, particularly infectious disease (e.g. the timely COVID-19 mRNA vaccines)^74–77^. However, there still remain several major hurdles that need to be overcome for mRNA-based therapeutics to be effective, many of which are directly linked to the intrinsic features of mRNA as a molecule. In particular, RNA is inherently unstable due its 2’-hydroxyl group, and degradation of the mRNA in solution by in-line nucleophilic attack, potentially aggravated by formulations, poses a major challenge^6^. Moreover, once delivered into patient cells, the candidate mRNA must outcompete other cytoplasmic mRNAs for the translation machinery and avoid the cellular mRNA degradation machinery to express maximum amounts of the desired encoded protein. Our study presents advances in tackling both of these challenges, building on two new high-throughput technologies.

Our first set of experimental findings derive from the integrated PERSIST-seq technology, which enables massively parallel evaluation of the effects of UTR and CDS sequence and structure on in-cell mRNA translation efficiency, in-cell mRNA stability, and in-solution stability. This technology identifies previously unknown tradeoffs and additive effects between different tunable aspects of mRNA performance as key determinants. Most important for optimizing these characteristics and their interdependencies, we find that in-cell mRNA stability may be a greater driver of protein output than high ribosome load particularly when proteins need to be expressed from a single dose of mRNAs for long periods of time. In particular, high translation efficiency mRNAs associated with the heaviest polysomes tend to be less stable: there is a ‘sweet spot’ for translation efficiency for maximal total protein output. This effect may derive from overcrowding of ribosomes on mRNA coding regions due to rapid initiation, which can induce sterical ribosome collisions that have recently been found to lead to translation-dependent mRNA decay^78,79^. Understanding this phenomenon is an exciting prospect for future mechanistic studies. Overall, we observed a wide range of UTR-dependent translation efficiencies in PERSIST-seq which could be further fine-tuned with specific UTR sequence or CDS structure alterations. We note that for applications with very short mRNAs (as are in use for multi-epitope cancer vaccines, and represented in our MEV sequences^80^) or much longer mRNAs (as are needed for antigens like the SARS-CoV-2 Spike protein), the best UTRs and the sweet spot for optimal translation efficiency may be different. In this respect, certain UTR combinations outperform *hHBB* in translation efficiency and include multiple cellular 5’ UTRs as well as, unexpectedly, the dengue virus 5’ and 3’ UTRs, which both individually increase ribosome loading, and combining them in one mRNA resulted in an additive effect. As illustrated with the SARS-CoV-2 5’ UTR, selective translational enhancers can be further identified through careful mutagenesis and deletion strategies aimed at narrowing selective regulatory regions within viral leader sequences. We also note that our experiments were limited to HEK293T as model human cells; the best UTRs for high protein output are likely to be cell-type dependent and therefore will vary from application to application. Our library of UTRs and the PERSIST-seq technology should be well-suited for discovering and leveraging the optimal UTRs for future applications with different protein targets and different cell types.

Our second set of experimental improvements relate to mRNA stability in aqueous solution. PERSIST-seq confirms predictions that extremely highly structured mRNAs can exhibit more than double the in-solution half-lives of conventionally designed mRNAs, with strong implications for improved storage in solution. Further design insights came from a new method called In-line-seq for high-throughput in-line probing^63,65^, applied to thousands of diverse short RNAs derived from the Eterna crowdsourcing platform. The In-line-seq results reveal a number of structural rules for mitigating in-solution RNA hydrolysis as well as simple sequence rules: a key determinant of in-solution RNA degradation is the presence of uridine and that RNA linkages 5’ of a uridine residue are particularly susceptible to degradation. Interestingly, this effect can be alleviated through inclusion of the nucleoside modification Ψ or m^1^Ψ. Synthesis of mRNAs with modifications – already in wide use for mitigating innate immune response and translational shutdown by mRNA therapeutics^67,68,74,75^ – is therefore a simple method to achieve significantly greater in-solution stability while sustaining protein expression.

Further leveraging our in-line hydrolysis data, we developed DegScore, a model for hydrolytic degradation that was independently validated on PERSIST-seq data and which enables *in silico* optimization of any RNA sequence. By combining optimal UTRs, DegScore optimization with RiboTree, and Ψ modification, we are able to achieve high mRNA stability and improved protein expression. We note that DegScore was trained on degradation from unmodified nucleotides that does not account for our observed stabilization via Ψ. Thus, future studies training similar models on degradation data from Ψ and other nucleoside modifications may result in improvements to CDS stabilization via algorithmic design. We also note that our study has focused on characterizing degradation of naked, non-formulated mRNAs in order to understand limitations on stability imposed by the fundamental biophysical and biochemical properties of RNA; such non-formulated mRNAs also appear optimal for certain applications including personalized cancer mRNA vaccines^81^. For applications that benefit from mRNA formulated in lipid nanoparticles or other carriers, we expect future studies applying PERSIST-seq and In-line-seq to formulated mRNA libraries to reveal additional powerful insights. Lastly, it has been proposed that highly structured mRNAs may retain their structure and in-solution stability under temperature shifts, mutations, and changes in UTRs, motivating the term ‘superfolder’ mRNAs^7^; it will be interesting to test these predictions through future PERSIST-seq studies.

Overall, we report a new mRNA design methodology that can dramatically enhance mRNA stability in aqueous solution while sustaining or even increasing protein expression inside cells. There does not have to be a tradeoff between mRNA structure, stability, and protein output. Looking ahead, our computational and experimental methods provide a platform to rapidly develop customized highly structured mRNAs for novel target proteins. As mRNA-based medicines are explored for a wide range of human diseases including cancer therapies, we hope that these insights and methods can help these medicines become more effective, manufactured at lower cost per patient, and more accessible and widely distributed to alleviate disease.

## SUPPLEMENTAL TEXT

### In-cell 5’ UTR sequence selection to determine rules of efficient translation initiation

Beyond comparing full UTR and CDS regions, we further sought to select for an optimally translating 5’ UTR sequence in an unbiased fashion from a complex sequence library (**Fig. S2**). Similar to sequence selection by enrichment through direct binding^82^ previously performed for mRNA-stabilizing 3’ UTR sequences^18^, we selected for highly translating transcripts by transfecting an mRNA reporter library with varying 5’ UTR sequences and harvesting mRNAs associated with heavy polysomes (**Fig. S2A**). We further enriched these libraries for highly translating transcripts over a total of five rounds of selection and re-transfection of the heavily ribosome loaded mRNAs from two independent starting pools (**Fig. S2A, B**). We compared them to input sequences of the initial and fifth selection round by RNA-seq. Using the *hHBB* 5’ UTR as our baseline (**Fig. S2A**), our 5’ UTR library design used the first 29 nt of the *hHBB* 5’ UTR followed by a 35 nt stretch of random sequences (N35, N=A,C,T,G) and the consensus Kozak sequence (GCCACC<u>AUG</u>G)^83,84^ upstream of the Nluc ORF.

First, we asked whether 5’ UTRs selected to be polysome-associated would increase the protein output compared to *hHBB*. We chose candidate 5’ UTRs in which we observed high read counts in the final round (≥15 reads), increasing representation across all selection rounds (FDR≤0.1), and >2-fold enrichment in the last round of selection compared to its input (**Fig. S2C**) and performed luciferase reporter assays with these mRNAs (**Fig. S2D**). Beside a wide range of luciferase activity driven by candidate 5’ UTR mRNAs, we surprisingly observed that none demonstrated luciferase activity that was significantly higher compared to *hHBB* 5’ UTR. Thus, although we are selecting for 5’ UTR reporter mRNAs of highest ribosome load, this unexpectedly decreases total protein output, which suggests the selected 5’ UTRs may have also impacted mRNA stability or translation elongation kinetics; a similar tradeoff is reported in the PERSIST-seq measurements described in the main text (**Fig. 2**).

To determine common features among the selected 5’ UTR sequences, we calculated position-specific short k-mer enrichment across the N35 region using *k*pLogo^85^ (**Fig. S2E**). We observed stronger enrichment/depletion of specific k-mers (165,611 k-mers tested in total) towards the 5’ and 3’ ends of the N35 stretch (**Fig. S2E**). In a confirmatory observation expected from ribosome scanning model of translation initiation, AUG triplets are significantly depleted across the N35 region (**Fig. S2F**). This effect is periodic and specific to two out-of-frame (frames 1 and 2) AUGs while in-frame AUG (frame 0) is not strongly affected, suggesting the negative impact of the competing upstream start codon except when it is in-frame to result in an N-terminally extended ORF and protein product. A variety of other interesting motifs are further observed, such as the depletion of guanine repeats (for example depletion of GGGG or GGG at the 3’ end of the N35, close to the fixed Kozak consensus) and uridine repeats throughout the 5’ UTR and enrichment of specific k-mers that suggest formation of short stem-loop structures promoting translation (**Table S3**). The latter is especially striking: for example, the 6-mer GUGAAC is strongly enriched towards the 5’ positions of the variable N35-mer region; GUGAAC is reverse complement to the last 6 nucleotides of the fixed HBB-29 region (GUUCAC), which would therefore be able to perfectly base-pair with each other and comprises an inverted repeat (**Fig. S1G**). The enrichment of the 6-mer peaks at the 4th to 6th nucleotide position downstream of the HBB-29 region, thus favoring an intervening length of 3 nt that would allow a 3-nt loop to form after base-pairing with the 6-mer stretches. Examining other possible inverted repeat k-mers in the variable region as 6-mer reverse complements sliding along the fixed region, we find that the stem may be formed up to around position −30 to the AUG. Such a pattern indicates that folding of a small stem-loop in the middle of the 5’ UTR under selection may actually be favored in mRNAs with heavy polysome load. This finding is in contrast to the typical expectations for secondary structures in 5’ UTRs to generally repress translation initiation. This finding is interesting because some synthetic small 5’ UTR RNA hairpins have previously been found to improve protein expression^86^. In sum, our sequence selection strategy formalizes previously predicted rules for 5’ UTR sequences that optimize ribosome load, and motivates an integrated approach to optimization of protein expression that jointly leverages our ribosome load dataset (**Fig. 1**) in parallel with our study of in-cell mRNA stability (**Fig. 2**).

---

**Figure S1.**
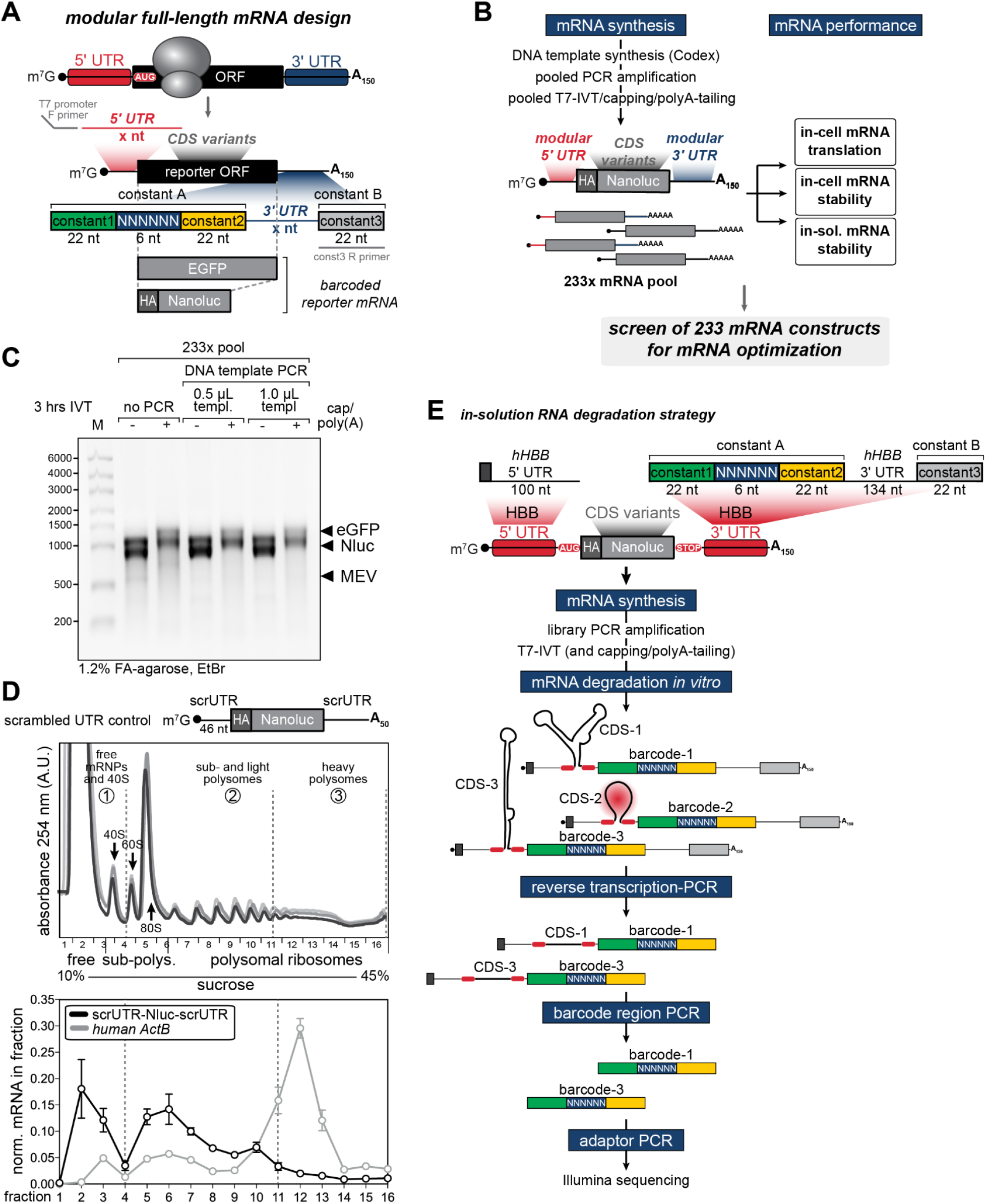
mRNA reporter design and in-cell and in-solution workflows with in-cell polysome validation. (A) Schematic for the 3’ UTR-barcoded mRNA reporter used to screen mRNA performance in a pooled format. The constant regions and barcode, which flank a variable 3’ UTR, were instrumental for amplifying and identifying hundreds of constructs simultaneously in each of the pooled experiments that comprise PERSIST-seq. The DNA templates for full-length mRNAs were synthesized on the Codex platform and amplified in a pooled PCR using primers complementary to the constant region (T7 promoter) preceding the variable 5’ UTR, and to the ‘constant3’ region following the variable 3’ UTR. (B) Summary of the workflow to progress from the individually synthesized DNA templates to the *in vitro* synthesized mRNA pool of 233 different constructs. We then use the same mRNA pool to screen mRNA performance in a three-pronged set of in-cell and in-solution expression and stability analyses. (C) Quality control of the 233-mRNA pool on a 1.2% formaldehyde (FA) gel stained with ethidium bromide (EtBr) after 3 hrs of *in vitro* transcription (IVT). The mRNA pool was analyzed before and after capping and polyadenylation. Pooled IVT is equally efficient with the starting template DNA pool with or without PCR-amplification of the DNA template pool. The three major bands corresponding to the three CDS types are indicated. The RiboRuler High Range RNA ladder (Thermo Fisher) is loaded for reference. (D) Polysome fractionation analysis of a transfected mRNA reporter. As an example, the distribution of an mRNA with short scrambled 5’ and 3’ UTRs 6 hrs after transfection into HEK293T cells was compared to the distribution of endogenous human *ActB* mRNA. RNA was extracted from fractions and quantified by qPCR with a RNA spike-in for normalization. Values are plotted as mRNA normalized per fraction. (E) In-solution RNA degradation strategy of barcoded mRNAs containing CDS variants with hHBB 5’ and 3’ UTRs. The differential degradation of CDS variants depends on their individual CDS structures. mRNA pools are degraded in solution by nucleophilic attack (red circle). After degradation, RT-PCR is performed to selectively amplify mRNAs that remain intact along their full length. Then, the barcode regions of these full-length mRNAs are PCR-amplified, adaptor-ligated, and prepared for Illumina sequencing.

**Figure S2.**
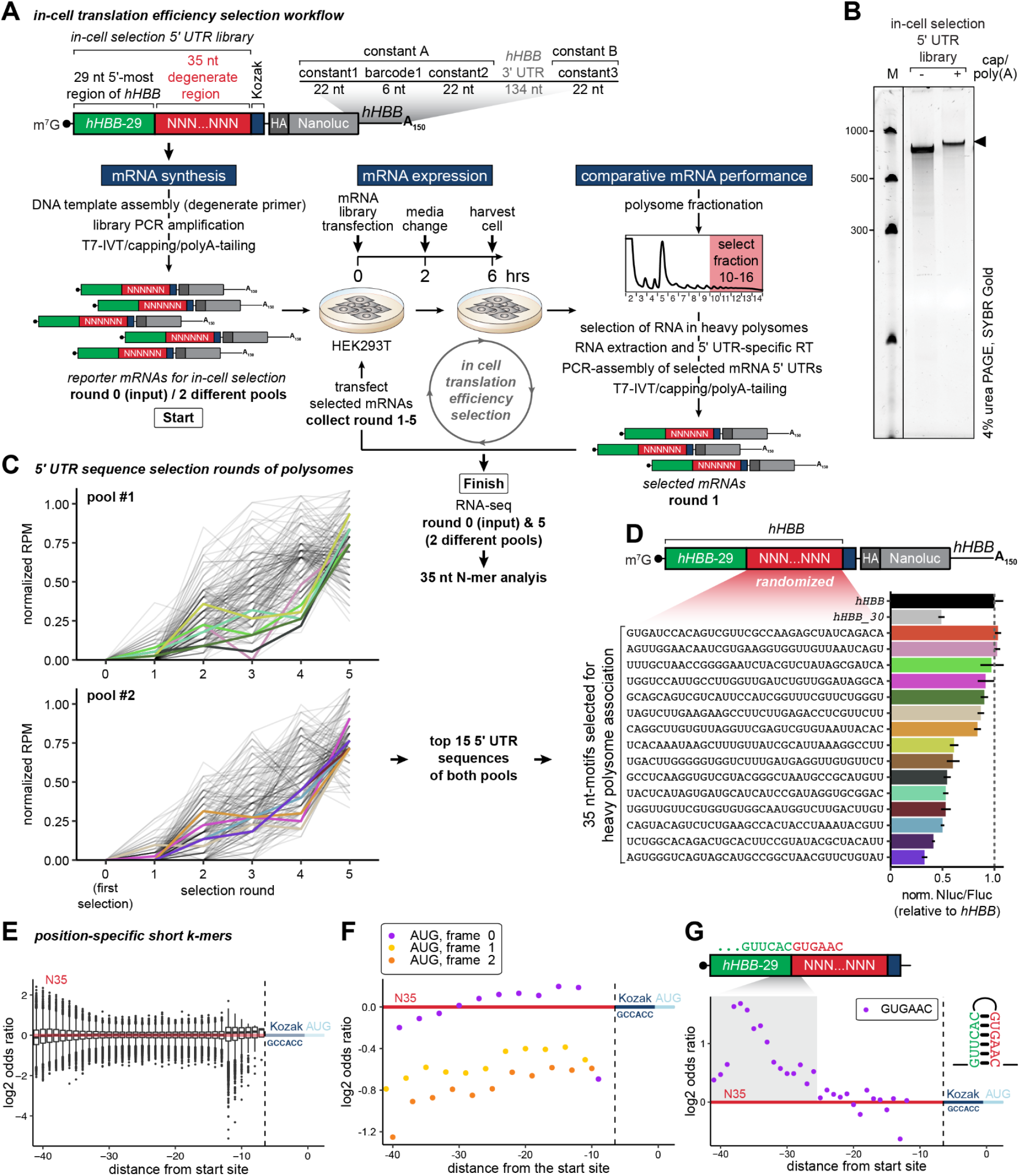
Sequential selection of high ribosome loaded mRNAs uncovers 5’ UTR sequences that contribute to protein abundance. (A) Overview of the in-cell selection assay designed to uncover 5’ UTR sequences that contribute to translational efficiency. First 29 nt of the human *HBB* 5’ UTR was chosen as the fixed 5’ region followed by the 35 nt long degenerate region and a constant Kozak consensus sequence (GCCACC). Selection of the variable 35 nt region is introduced by subsequent re-transfection of only the mRNAs purified from the heavy polysomal fractions. (B) Denaturing urea-polyacrylamide gel for the quality control of the *in vitro* transcribed mRNA 5’ UTR selection library before and after the 5’ cap and polyA-tail, shown for the selection round 0. All consecutive selection rounds yielded similar libraries. The Low Range ssRNA Ladder (NEB) was loaded for reference. The gel was stained with SYBR Gold (Thermo Fisher). (C) Normalized reads per million (RPM) of the top 5’ UTR sequences (FDR≤0.1) over the course of the selection rounds. Colored lines indicate mRNAs that were chosen for luciferase reporter assays (15 in total from two independent starting pools; ≥15 final round read count, ≥2-fold final round enrichment over input). (D) Normalized Nluc/Fluc luciferase activities of the top 15 mRNAs from (C). The 35-nt variable region in the 5’ UTR of the polysome selected mRNAs are listed along the y-axis. Their luciferase activity is plotted on the x-axis relative to *hHBB*. HBB-29 contains only the first 29 nt of the hHBB 5’ UTR. (E) Boxplot of all log2 odds ratios of k-mers (2≤k≤6) between the final polysome selection round and the initial starting pool. Higher variations are observed towards either 5’/3’ ends of the 35 nt variable region. Most of the significant k-mers in the 3’ positions are depletions. (F) Depletion of out-of-frame (+1- and +2-frames) AUGs within the 35 nt variable region following the polysome selection rounds. In-frame AUGs (0-frame) are weakly depleted or even show minor enrichment closer to the 3’ end. (G) Enrichment of the 6-mer motif GUGAAC following polysome selection. GUGAAC is reverse complementary to the 3’ end of the fixed 29-nt region of the 5’ UTR (GUUCAC). The enrichment towards the 5’ end of the variable region and its peak at the 4th to 6th nucleotides downstream of the end of the fixed region may indicate favorability of small stem loop structure for increased ribosome loading.

**Figure S3.**
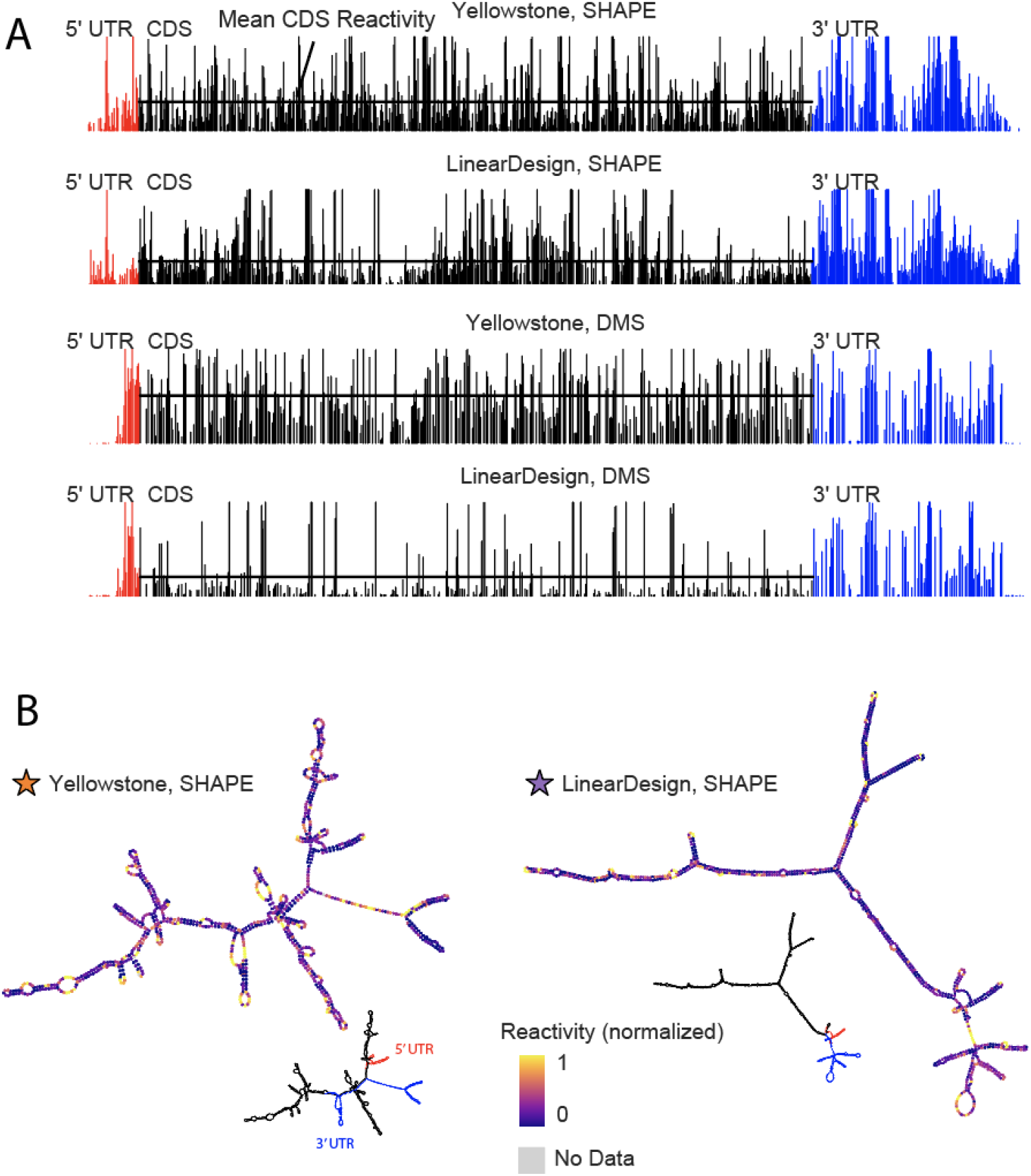
Chemical structure probing of Yellowstone and LinearDesign-1 RNAs. (A) SHAPE and DMS reactivity per sequence position of Yellowstone and LinearDesign-1. (B) MFE structures derived using SHAPE reactivity.

**Figure S4.**
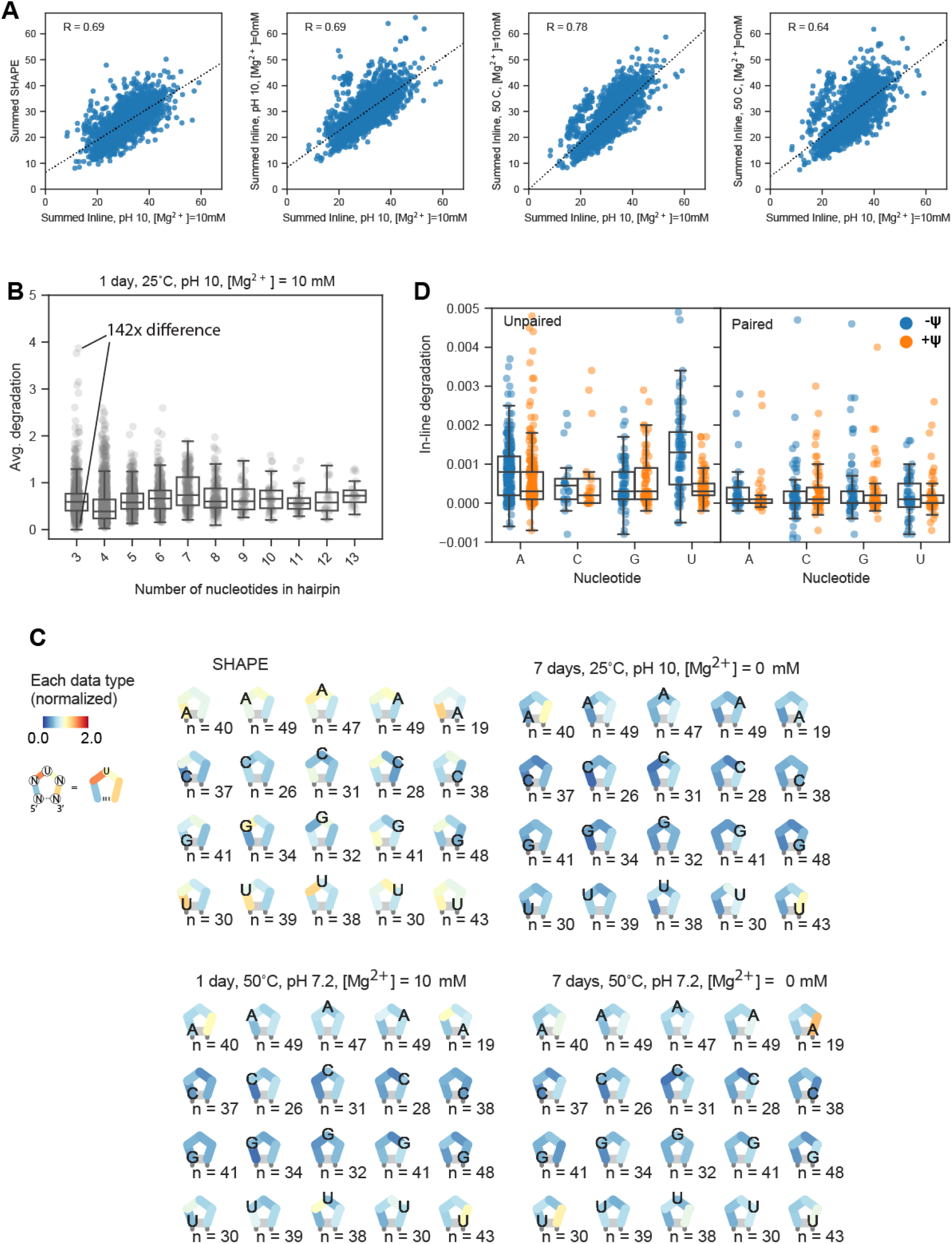
Features of RNA degradation as determined by In-line-seq. (A) Correlation of summed in-line degradation per construct at pH 10, 25°C, [Mg^2+^] = 10 mM, 1 day conditions, to SHAPE reactivity and other in-line degradation conditions tested. (B) Dynamic range of average reactivity for hairpin loop degradation for in-line degradation at pH 10, 25°C, [Mg^2+^] = 10 mM, 1 day conditions. (C) Sequence/location dependency of triloop reactivity and degradation for other three experimental in-line degradation conditions tested (see **Fig. 3C**). (D) In-line degradation for 8 constructs measured one-by-one with capillary electrophoresis, in absence and presence of pseudouridine. Left panel depicts nucleotides predicted to be unpaired, right panel depicts nucleotides predicted to be paired in ViennaRNA structure.

**Figure S5.**
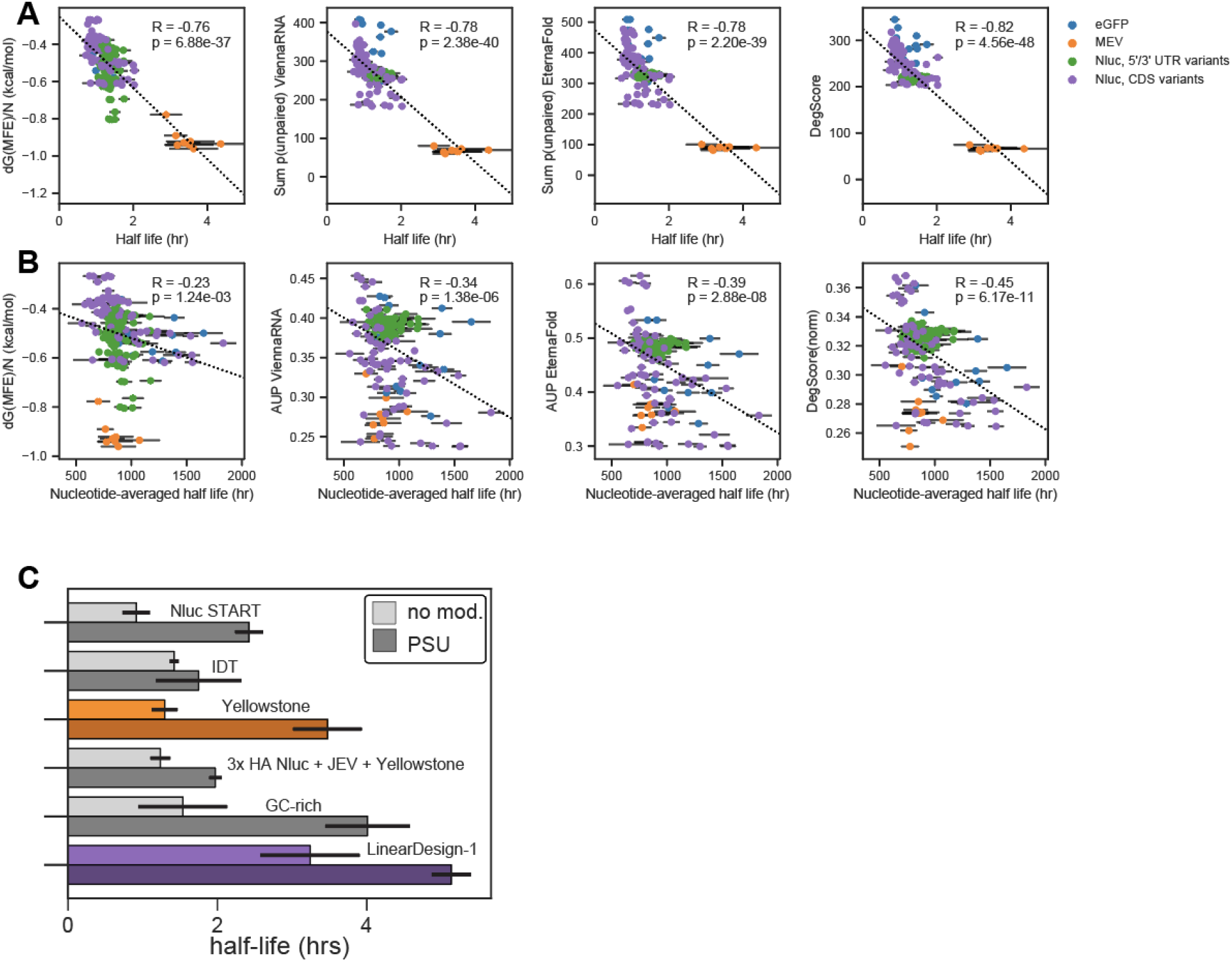
Correlation between the 233-mRNA pool in-solution half-life and predictors for RNA degradation. (A) Correlation between in-vitro half-lives and dG(MFE), Sum p(unpaired) calculated in ViennaRNA and EternaFold, and DegScore across all model mRNA types tested. (B) Correlation between in-vitro half lives, normalized to RNA length, and dG(MFE), Average p(unpaired) (AUP) in ViennaRNA and EternaFold, and DegScore across the Nanoluciferase and eGFP constructs. (C) One-by-one characterization of in-vitro half-lives of 6 model mRNAs, characterized with U and with pseudouridine.

**Figure S6.**
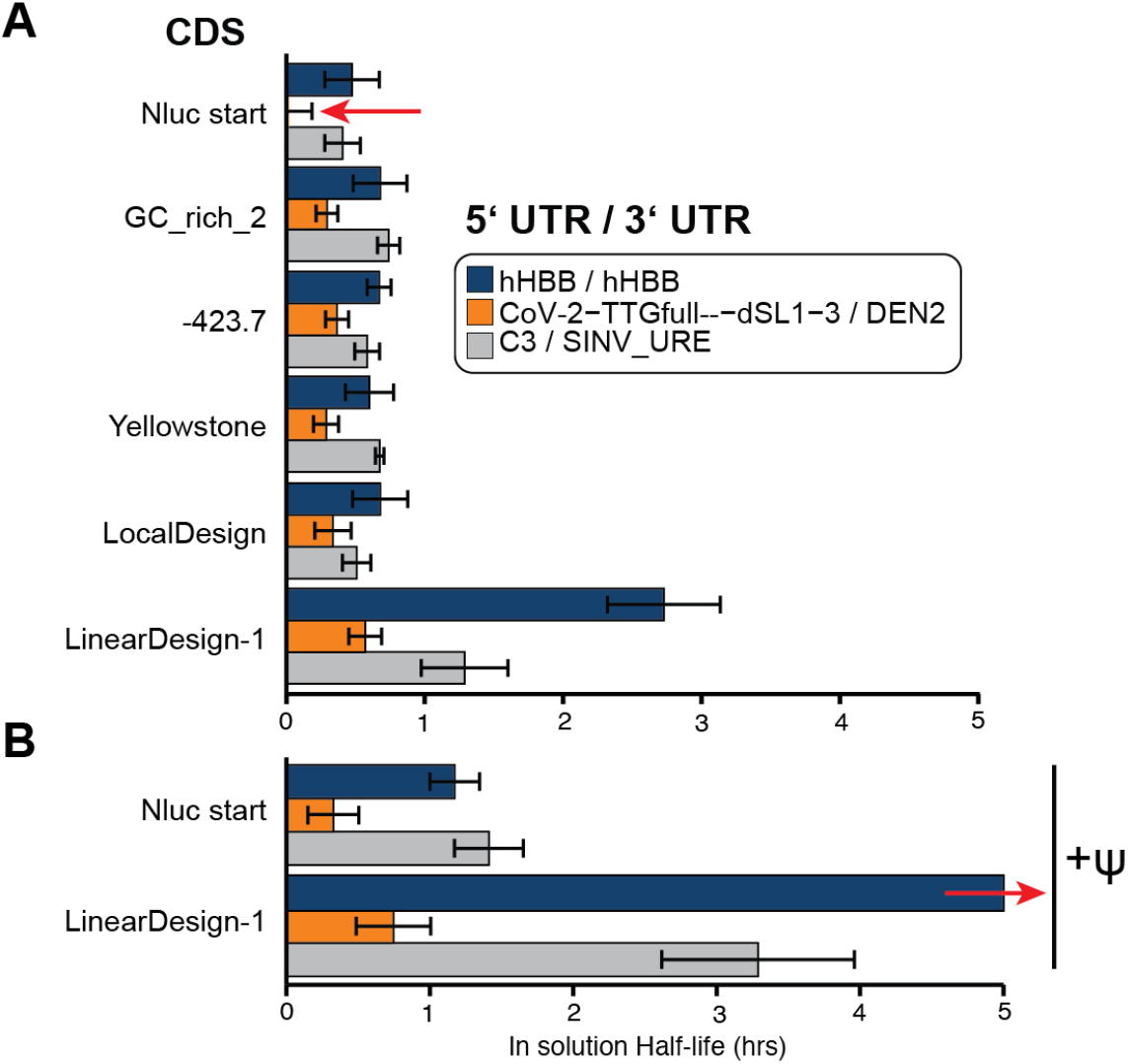
Effect of UTR and modified nucleosides on in-solution half-life. (A) 6 select CDS designs were combined with three different pairs of 5’ and 3’ UTRs and the in-solution half-lives were measured. The half-life of ‘Nluc start’ with CoV-2-UUG-UUGfull-dSL1-3/DEN2 UTRs (red arrow) could not be accurately measured as it was outside the dynamic range of the experiment; data represent an upper bound. (B) Two model RNAs from Panel A were synthesized with pseudouridine and in-solution half-lives were measured. The half-life of “LinearDesign-1” with hHBB/hHBB UTRs containing pseudouridine (red arrow) was not accurately captured as this RNA persisted beyond the range of the experiment; data reflect an approximate upper bound. Error bars indicate standard deviations from 1000 bootstrapped exponential fits for half-life calculation

**Figure S7.**
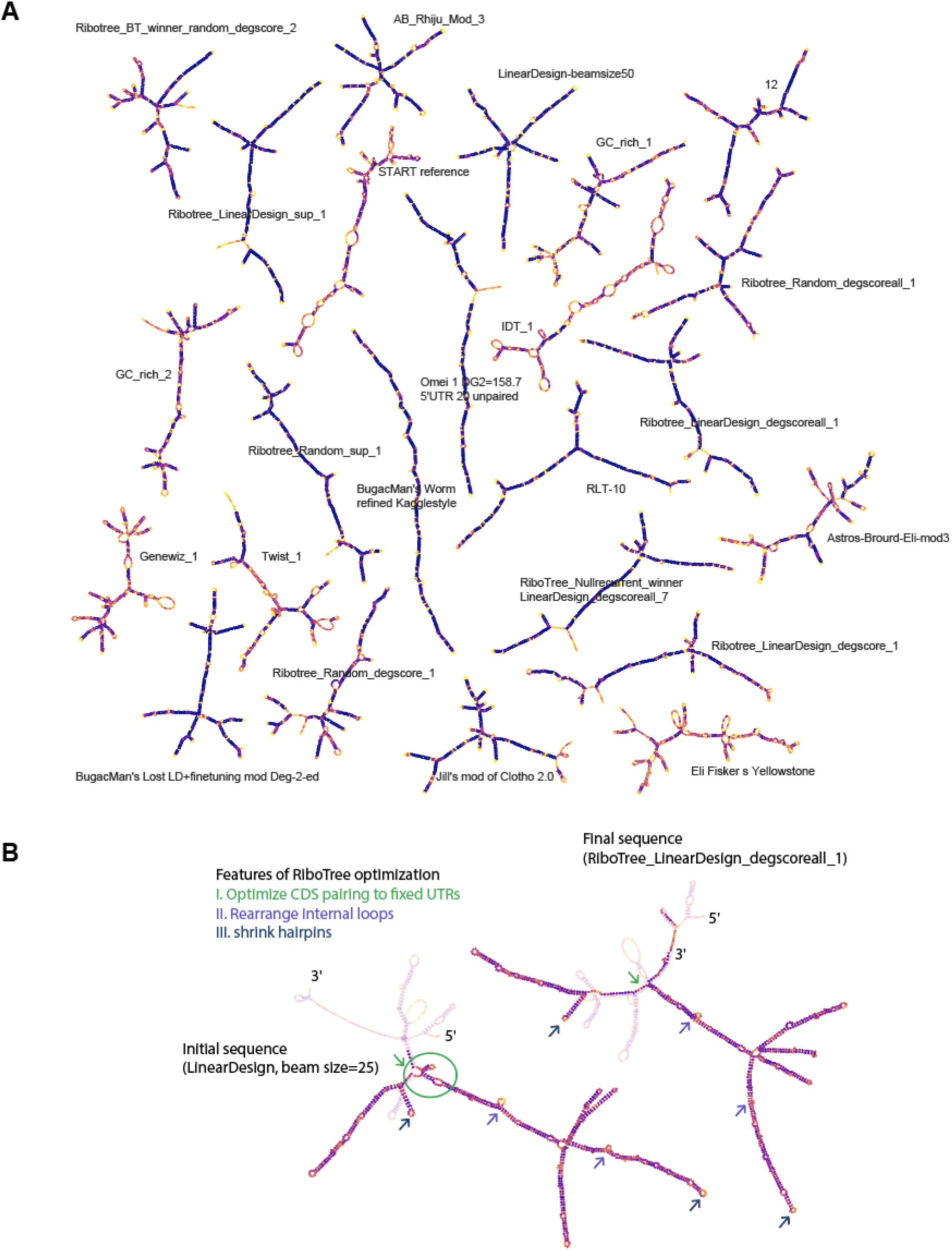
Overview of predicted RNA secondary structures of constructs in Fig. 4C. (A) Predicted secondary structures of final tested Nanoluciferase constructs, colored by AUP. (B) Comparison of secondary structure of starting sequence used for RiboTree_LinearDesign_degscoreall_1 and annotations of changes.

**Figure S8.**
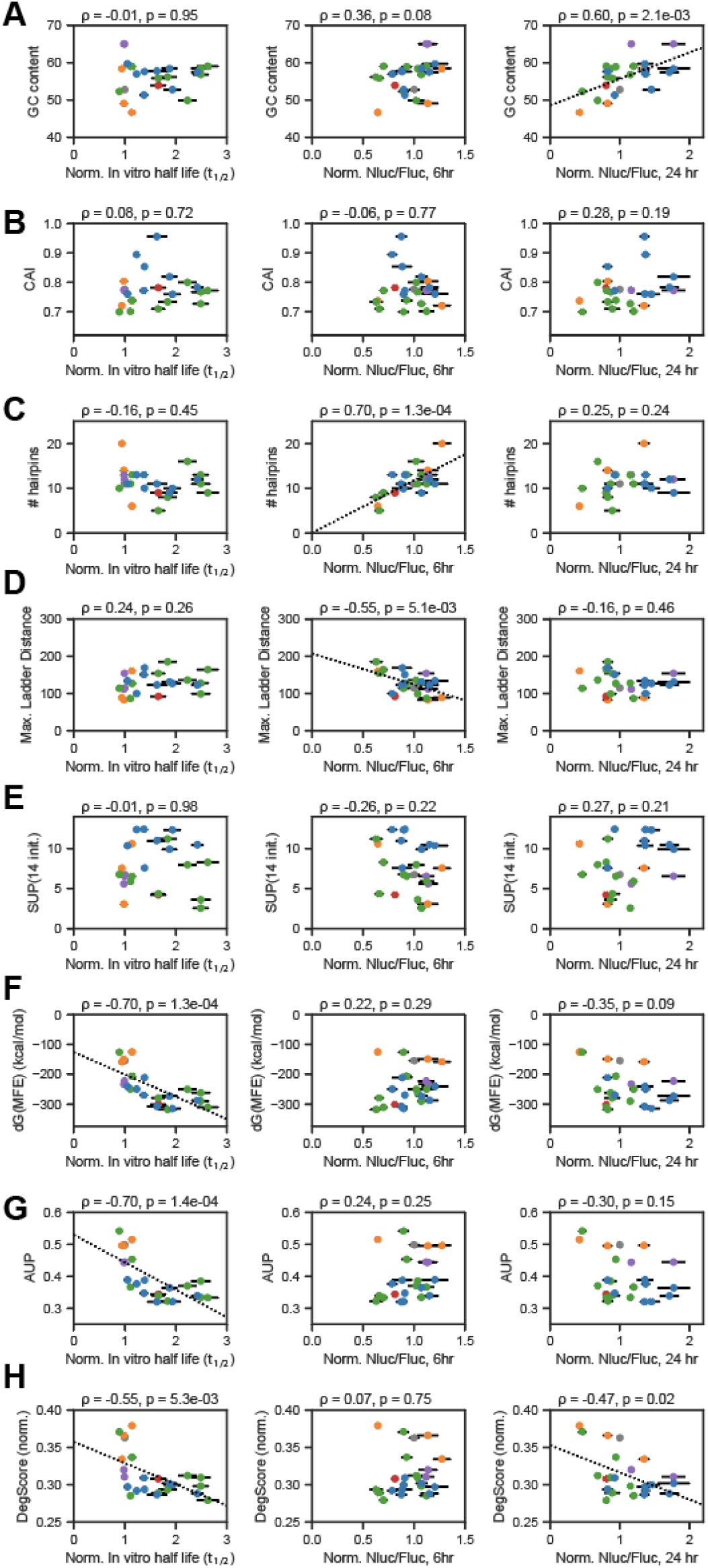
Correlation of experimental data for 24 Nluc constructs to predicted degradation and structure metrics. (A) Correlation of normalized in-solution half-life, normalized Nluc expression at 6 hours and 24 hours, tested for correlation to (A) GC content; (B) CAI; (C) number of hairpins; (D) Maximum ladder distance, the maximum length of contiguous helices in the secondary structure; (E) SUP(14 init.), the summed unpaired probability of the first 14 nucleotides; (F) dG(MFE); (G), AUP (average unpaired probability); (H) DegSore. Notably, AUP and dG(MFE) have higher correlation to in-solution half-lives than DegScore; this is possibly because the DegScore model was not trained with data on pseudouridine.

## ACKNOWLEDGEMENTS

We would like to thank the Barna and Das lab members for support and constructive criticism and reading of and helpful comments on the work, and Eterna Players for providing sequences and for critical reading of the manuscript. Eterna participant usernames and full names, when volunteered, are provided in **Table S7**. We thank Ramya Rangan for advice on SARS-CoV-2 structure. We thank Ivan N. Zheludev for performing pilot experiments. We are grateful to Kelsey L. Hickey and Jonathan S. Weissman (UCSF, San Francisco, CA, USA); and Conor J. Howard (UCSF, San Francisco, CA, USA) for kindly sharing plasmids. We would like to thank John Coller and Dhananjay Wagh of the Stanford Functional Genomics Facility (SFGF), and Michael Eckart, Kyle Fukui and Jennifer Okamoto (Protein and Nucleic Acids Facility) for experimental support. We thank Liang Huang (Oregon State University, Baidu Research USA) for discussions of LinearDesign. RiboTree calculations were performed on the Stanford Sherlock cluster. We thank Camilla Kao, Howard Y. Chang (Stanford University); Hani Choudhry (King Abdullaziz University); Anthony Goldbloom, Maggie Demkin, and Walter Reade (Kaggle, Inc.); and Dani Braun, Austin Chen, Sharif Ezzat, Abhi Garg, Johannes Häggqvist, Tamas Kalman, Kevin Lin, and Amine Rehioui for development and advice important for launching online components of the OpenVaccine challenges. We thank Helga Leppek for providing the Barna lab with handmade masks at the onset of the pandemic for essential lab work and Lisa Sharp and Jessica Corkern (Stanford Biochemistry Department) for supporting Das laboratory experiments. This work was supported by NIH grants R01HD086634 (M.B.) and R35 GM122579 (R.D.), the National Science Foundation GRFP (H.K.W.S., D.S.K.), a Benchmark Stanford Graduate Fellowship (G.W.B, C.A.C.), a Stanford ChEM-H “Stanford RISE” seed grant (H.K.W.S.), a Canadian Institutes of Health Research Postdoctoral Fellowship (C.K.), Paul and Daisy Soros Fellowships for New Americans (G.C.T), Stanford Medical Scientist Training Program (A.X., G.C.T.), and a Human Frontier Science Program Fellowship (K.F.). D.R. is supported by an EMBO Long-term Fellowship (ALTF 1042-2019) and a Human Frontier Science Program Fellowship (LT000218/2020-L). A.X. is supported by a Stanford Bio-X/SIGF fellowship and NIH fellowship 1F30HD100123. K.L. is supported by an EMBO Long-Term Fellowship (ALTF 539-2015), is the Layton Family Fellow of the Damon Runyon Cancer Research Foundation (DRG-2237-15), and is supported by the Katharine McCormick Advanced Postdoctoral Scholar Fellowship to Support Women in Academic Medicine (2019). M.B. is a New York Stem Cell Foundation Robertson Investigator. Further financial support came from gifts to the Eterna OpenVaccine project from donors listed in **Table S8**.

## METHODS

### *In vitro* transcription of reporter mRNAs

Preparation of mRNAs were based on *in vitro* transcription from DNA templates. DNA templates were amplified by PCR using AccuPrime Pfx (Life Technologies, 12344024) and purified using the Monarch PCR & DNA Cleanup Kit (NEB, T1030L). The source of the 3xHA-Nluc starting CDS (“Nluc start”) is derived from the pcDNA3.1-5’UTR-3xHA-Nluc plasmid encoding the HA-tagged Nanoluc CDS^43^. Individual template DNA or the 233-mRNA library was amplified from linear DNA synthesized on a BioXP 3200 system (Codex DNA) or by Twist Bioscience, using the fixed forward (T7_F_28nt) and reverse (const3_R) primer. The forward primer binds to the T7 RNA polymerase promoter common in DNA template for all mRNA designs; the reverse primer is complementary to a common “const3” region at the end of all tested mRNA 3’ UTRs. For the IVT template pool, individual DNA templates were pooled for a template pool of hundreds of constructs at an equimolar concentration and are amplified with outer primers in a pooled format. For the pooled template, 1 μL of each construct (~20 ng/μL stock concentration) was pooled to be used as the PCR template. The Pfx PCR contained the following: 2.5 μL 10x Pfx buffer, 0.25 μL forward primer (100 uM), 0.25 μL reverse primer (100 uM), 0.75 μL DMSO (NEB), 0.25 μL Pfx Polymerase (Thermo), 20.5 water, and 0.5 μL template DNA (~20-50 ng/ul), in a total 25 μL reaction with the following program: 2 min at 95°C; 10 sec at 95°C; 30 sec at 58°C; 30s or 1 min at 68°C; cycled 9x; final extension of 5 min at 68°C. PCR reactions were purified with Monarch PCR & DNA Cleanup Kit (NEB, T1030L). For the hHBB-Fluc control mRNA, the DNA template was amplified from the pGL3-HBB plasmid^87^ using the primers KL588/KL589 which yielded a PCR product of 1,750 kb in length. For cloning the MALAT1 ENE 3’ UTR stem-loop, we first amplified the ENE region using primers ENE-1/ENE-2 with flanking constant regions. The resulting amplicon was assembled with a hHBB-Nluc sequence that lacked a 3’ UTR but maintained a unique barcode using a NEBuilder HiFi Assembly Kit (NEB, ES2621).

*In vitro* transcription was performed with the MEGAscript T7 kit (Ambion, AM1333) according to the manufacturer’s instructions. A 20 μL transcription reaction contained max. 5 μg linear DNA template, 4 mM of each NTP (Ambion), 2 μL/ 200 U MEGAscript T7 RNA polymerase (Ambion) and 1x T7 MEGAscript Transcription Buffer (Ambion). After a total incubation for 3 hours at 37°C, the DNA was digested by addition of 1 μL/2 U Turbo DNase (Ambion, AM2238) for 15 min at 37°C. For pseudouridylated mRNAs, pseudouridine triphosphate (Trilink Biotechnologies, N1019-5) was substituted for uridine triphosphate at an equivalent concentration. mRNA was purified using MegaClear columns (Thermo Scientific, Ambion, AM1908). A 20 μL reaction usually yielded 100-150 μg of RNA.

For mRNA transfection of HEK293T cells, m^7^G-capped and polyadenylated mRNAs were generated as follows. In vitro transcribed mRNA was then m^7^G-capped and polyadenylated using the ScriptCap m7G Capping System (CellScript, C-SCCE0625) and A-Plus Poly(A) Polymerase Tailing Kit (CellScript, C-PAP5104H), respectively, according to the manufacturer’s instruction with the following modifications. Aliquots of 30 μg of each RNA were processed in parallel, diluted to 34.25 μL in water and heated for 5 min at 65°C to denature and placed on ice. The 50 μL capping reaction contained 5 μL 10x ScriptCap buffer (Cellscript), 5 μL 10 mM GTP (Cellscript), 2.5 μL 2 mM S-adenosyl-methionine (SAM, 20 mM stock, Cellscript), 1.25 μL ScriptGuard RNase Inhibitor (Cellscript), and 2 μL Capping enzyme (20 U, Cellscript, 10 U/μL). For the capping step, the 37°C incubation was performed for 1 hour and the capped RNA was placed on ice. Polyadenylation was performed from the resulting RNAs without purification in between. The polyA reaction contained 30 μg of capped mRNA in 50 μL, 6.6 μL 10x A-Plus polyA tailing buffer (Cellscript), 6.6 μL 10 mM ATP (Cellscript), 0.3 μL ScriptGuard RNase Inhibitor (Cellscript), and 2.5 μL A-Plus PolyA Polymerase (10 U, 4 U/μL, Cellscript) in a total reaction volume of 66 μL. We aimed to add a 150 nt-long polyA-tail for which we incubated the capped mRNA for 30 min at 37°C with 10 U of polyA enzyme, after which the reaction was placed on ice. The mRNA was again purified using MegaClear columns. mRNA concentration was determined on a Nanodrop 2000 (Thermo Fisher). This usually yields 30-40 μg of capped and polyadenylated mRNA. mRNA quality was determined by 4% urea-PAGE, 1% formaldehyde agarose gel or capillary electrophoresis with a Agilent 2100 Bioanalyzer (Agilent Technologies). A list of all primer sequences used are provided in **Table S6**.

### Cell culture and transfection

HEK293T (ATCC: CRL-3216) cells were cultured in Dulbecco’s Modified Eagle’s Medium (DMEM, Gibco, 11965–118) containing 2 mM L-glutamine, supplemented with 10% fetal bovine serum (EMD Millipore, TMS-013-B), 100 U/ml penicillin and 0.1 mg/ml streptomycin (EmbryoMax ES Cell Qualified Penicillin-Streptomycin Solution 100X; EMD Millipore, TMS-AB2-C or Gibco, 15140–122) at 37°C in 5% CO_2_-buffered incubators. For transfection of pooled 5’ m^7^G-capped and poly(A)-tailed RNAs, 5.0 × 10^6^ HEK293T cells were seeded in a 10 cm plate 24 h before transfection. 10 μg of pooled RNAs were transfected using Lipofectamine MessengerMax as per manufacturer’s instructions (Life Technologies). Media was changed 3 h after transfection and replaced with complete DMEM supplemented with 10% FBS and Pen/Strep. For transfections of individual m^7^G-capped RNAs, 3.0 × 10^4^ HEK293T cells were seeded per well 24 h before transfection in a 96-well plate. Subsequently, 10 ng of Nluc RNA was co-transfected with 20 ng of m^7^G-capped HBB-Fluc control RNA using Lipofectamine MessengerMax as per manufacturer’s instructions (Life Technologies). A list of all primer sequences used are provided in **Table S6**. All oligonucleotides were purchased from IDT.

### Sucrose gradient fractionation analysis

Cell culture media was replaced with cycloheximide (MilliporeSigma, C7698-1G) containing media at 100ug/mL. After 2 minutes, cells were washed, trypsinized and harvested using PBS, trypsin, and culture media containing 100 g/mL cycloheximide. ~10×10^6^ cells were resuspended in 400 μL of following lysis buffer on ice for 30 min, vortexing every 10 min: 25 mM Tris-HCl pH 7.5, 150 mM NaCl, 15 mM MgCl2, 1 mM DTT, 8% glycerol, 1% Triton X-100, 100 μg/mL cycloheximide, 0.2 U/μL Superase-In RNase inhibitor (ThermoFisher Scientific, AM2694), 1x Halt protease inhibitor cocktail (ThermoFisher Scientific, 78430), 0.02 U/μL TURBO DNase (ThermoFisher Scientific, AM2238). After lysis, nuclei were removed by two step centrifuging, first at 1300 g for 5min and second at 10000 g for 5min, taking the supernatants from each. 25%-50% sucrose gradient was prepared in 13.2mL ultracentrifuge tubes (Beckman Coulter, 331372) using Biocomp Gradient Master with the following recipe: 25 or 50% sucrose (w/v), 25 mM Tris-HCl pH 7.5, 150 mM NaCl, 15 mM MgCl_2_, 1 mM DTT, 100 μg/mL cycloheximide. The lysate was layered onto the sucrose gradient and ultracentrifuged on Beckman Coulter SW-41Ti rotor at 40000rpm for 150min at 4°C. The gradient was density fractionated using Brandel BR-188 into 16×750 μL fractions, and in vitro transcribed spike-in RNA mix (120002B1, 120010B1, 220023B1, 310333T3; 1000, 100, 10, 1-fold dilutions respectively) were added to each fraction. 700 μL of each fraction was mixed with 100μL 10% SDS, 200μL 1.5M sodium acetate, and 900μL acid phenol-chloroform, pH 4.5 (ThermoFisher Scientific, AM9720), heated at 65°C for 5min, and centrifuged at 20000g for 15min at 4°C for phase separation. 600μL aqueous phase was mixed with 600 μL 100% ethanol and RNA was purified on silica columns (Zymo, R1013).

### Luciferase activity assay after mRNA transfection

Media from transiently transfected HEK293T cells was aspirated and cells were lysed in 40 μL of 1x passive lysis buffer from the Dual-Luciferase Reporter Assay System (Promega, E1980) and either directly assayed or frozen at −20°C. After thawing, 20 μL of supernatant was transferred to a new plate and assayed for luciferase activity using the Nano-Glo Dual-Luciferase Reporter Assay System (Promega, N1610) to measure Firefly (Fluc) and NanoLuc (Nluc) luciferase activities. In particular, 50 μL of ONE-Glo Ex Reagent was added to each well of lysate and incubated for 3 minutes at room temperature before measuring Fluc activities. Subsequently, 50 μL of NanoDLR Stop & Glo reagent was added to each well, and incubated for 10 min at room temperature before measuring luciferase activities on a GloMax-Multi (Promega) plate reader. Luciferase reporter activity is expressed as a ratio between Nluc and Fluc. Each experiment was performed a minimum of three independent times. Because this assay relies on accumulation of luciferase in the cytosol, any signal peptides sequences (Table S1) were removed from the CDS for templates and mRNA for these transfection and luciferase activity experiments.

### Quantitative RT-PCR (RT-qPCR) Analysis

RNA-transfected HEK293T cells were first lysed and separated by sucrose density gradient fractionation as described above. From each fraction. RNA was purified by acidic phenol/chloroform followed by isopropanol precipitation. 0.5 mg of RNA was converted to cDNA using iScript Supermix (Bio-Rad, 1708840). cDNA was synthesized from 100-200 ng of total RNA using iScript Supermix (Bio-Rad, 1708840) containing random hexamer primers, according to the manufacturer’s instructions. PCR reactions were assembled in 384-well plates using 2.5 μL of a 1:4-1:5 dilution of a cDNA reaction, 300 nM of target-specific primer mix and the SsoAdvanced SYBR Green supermix (Bio-Rad, 1725270) in a final volume of 10 μL per well. Data were analysed and converted to relative RNA quantity using CFX manager (BioRad). For sucrose gradient fractions, the amount of RNA from individual fractions was expressed as a fraction of the total RNA collected from all fractions. Primers were used at 250 nM per reaction. A list of all primer sequences used for qPCR are provided in **Table S6**.

### In cell and in-solution RNA degradation time courses

For in-cell RNA stability, the 233-member *in vitro* transcribed mRNA pool (m^7^G-capped and polyA) was transfected into HEK293T cells as described above and RNA was harvested at 1, 7, 12, and 24 h in Trizol (ThermoFisher Scientific, 15596026). RNA was extracted from the aqueous phase on silica columns (Zymo, R1013).

For in-solution RNA degradation experiments, 750 ng of the 233-mRNA pool (not m^7^G-capped or polyA) was incubated in 30 μL of Degradation Buffer (50 mM CHES at pH 10 and 10 mM MgCl_2_) and collected over 10 time points: 0, 0.5, 1, 2, 3, 4, 5, 6, 16 and 24 h. To each sample, 15 μL of 0.5 M Tris-HCl pH 7 and 3 μL of 0.5 M EDTA-Na was added to quench the degradation. The integrity of each sample was checked by loading 5 μL of total RNA alongside a spike-in control (P4P62HP, 50 ng) onto a PAGE-Urea-TBE gel and visualized by SYBR Gold (Thermo Fisher). Subsequently, RNA was purified using Ampure beads + 40% polyethylene glycol 8000 (7:3) and checked again by PAGE-Urea-TBE gel and visualized by SYBR Gold.

### Library preparation and amplicon Sequencing

Up to 250 ng RNA in 2.75 μL was mixed with 0.25 μL 2 μM RT_Const2_N12_Read1Partial (**Table S6**) and 0.25 μL 10mM dNTPs each. The RNA samples were then denatured at 65°C for 5 min and chilled to 4°C. 1.75 μL reverse transcription mix was added to 5 μL total reaction volume: 1 μL 5x Superscript IV buffer, 0.25 μL 10mM DTT, 0.25 μL Superase-In (ThermoFisher Scientific, AM2694), 0.25 μL Superscript IV (Thermo 18091050). The reaction was incubated at 55°C for 45 min and inactivated at 80°C for 10 min.

First round PCR was performed under following conditions: 1 μL RT reaction, 10 μL 2x Q5 Hot Start Master Mix (NEB M0494S), 0.2 μL 100x SYBR (Thermo S7563), 1 μL 10uM Read1Partial_F, 1 μL 10 uM 50:50 Hbb_Fwd:Nluc_Fwd mix in 20 μL total volume. Cycling conditions were: 98°C for 60 sec, and 15 cycles of 98°C for 10 sec, 68°C for 10 sec and 72°C. Second round PCR was performed under the following conditions: 1 μL first round PCR, 10 μL 2x Q5 Hot Start Master Mix, 0.2 μL 100x SYBR, 1 μL 10 uM Read1Partial_F, 1 μL 10 uM Read2Partial_Const1_R in 20 μL total volume. Cycling conditions were: 98°C for 60 sec, and 5 cycles of 98°C for 10 sec, 72°C for 5 sec. Sequencing adaptors were added using the following conditions for final round PCR: 1 μL second round PCR, 10 μL 2x Q5 Hot Start Master Mix, 0.2 μL 100x SYBR, 1 μL 10 μM NEBNext Index Primer (NEB E7335, NEB E7500, NEB E7710, NEB E7730, NEB E6609), 1 μL 10 μM NEBNext Universal PCR Primer in 20 μL total volume. Cycling conditions were: 98°C for 60 sec, and 5 cycles of 98°C for 10 sec, 72°C for 5 sec. All barcoded samples were then pooled at equal volumes and purified with 1.1x SPRIselect beads (Beckman Coulter B23317). Sequencing was performed at the Stanford Functional Genomics Facility (SFGF) at Stanford University, on an Illumina NextSeq 550 instrument, using a high output kit, 1×76 cycles. Primer sequences and the sequencing construct layout are provided in **Table S6**.

### Amplicon sequencing data analysis

After bcl conversion and demultiplexing with Illumina bcl2fastq, the constant regions were trimmed using cutadapt^88^. The trimmed reads were aligned to the indexed reference of barcode sequences using Bowtie2 with the following options: -L 11 -N 0 --nofw ^89^. The alignments were deduplicated based on UMIs using UMIcollapse^90^ with -p 0.05 and counted using samtools idxstats. This pipeline yields a matrix of barcode read counts where rows are the different constructs in the library and columns are the different samples.

The count matrix was log transformed and normalized column-wise using a linear fit on the dilution series of spike-in constructs in each sample. For the calculation of RNA degradation coefficients in cells, we carried out a linear fit to log RNA abundance from the time course data, i.e. we fit an expression of *Y* = β_0_+β_1_*t* where *Y* is the normalized log RNA abundance and *t* is the number of hours after transfection; β_1_ is the degradation constant. For the calculation of in solution degradation coefficients, sufficient data points were available to carry out a nonlinear fit directly to an exponential model, i.e. we fit an expression of *y* = *A* exp(−**τ**/*t*), where y is the fraction intact (RNA abundance normalized to initial abundance), A is the amplitude, *t* is the time of incubation in degradation buffer in hours, and **τ** is the degradation time constant.

For polysome profiles, percent RNA abundances for each fraction were first calculated by scaling per-fraction values by the sum of all fractions. For the heatmap displays in the figures, column medians were also subtracted from each percent RNA value. For the calculation of ribosome load, the matrix of percent RNA abundances in fractions 4-9 (1-3 are free RNP fractions, and >9 have negligible abundance) were first multiplied by a weight vector representing the number of ribosomes in each fraction as determined by the A260 trace from the fractionator, then the weighted abundances were summed across the row. For the calculation of polysome to monosome ratio, the sum of fractions 7-9 (>3 ribosomes) abundances were divided by fraction 4 (80S) abundance. For the calculation of monosome to 40S/60S ratio, fraction 4 (80S) abundance was divided by the sum of fraction 2 (40S/60S) abundance.

To calculated the expected protein levels assuming first order kinetics of mRNA translation and mRNA/protein decay, the following differential equations are used:

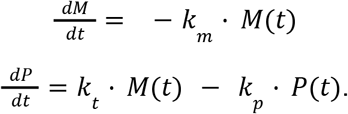

where *dM/dt* and *dP/dt* are rates of change in mRNA and protein levels, respectively; *M(t)* and *P(t)* are moles of mRNA and protein at time *t*, respectively; *k_t_* is the translation rate constant; and *k_m_* and *k_p_* are rate constants of mRNA and protein decay, respectively. The analytical solution for *P(t)* is proportional to:

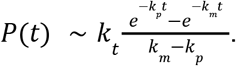

where *m_0_* is the mass of mRNA present at *t=0*, and *l* is the mRNA length in nucleotides. *k_p_* is set to 0 since Nluc protein has negligible degradation as measured by luciferase activity in transiently Nluc-expressing HEK293 cells for at least 6 hours after cycloheximide treatment, which allows assessment of protein degradation in the absence of further translation^91^. *k_m_* is the degradation constant obtained from the linear fit of in-cell time course RNA data (−β_1_ above). *k_t_* is the ribosome load calculated by summing weighted RNA abundances from polysome profile data.

### SHAPE and DMS chemical mapping of full-length mRNAs

For DMS-based chemical mapping of the LinearDesign-1 and Yellowstone mRNAs, 1 μg of RNA was brought to 10 μL in water, unfolded at 95 °C for 2 minutes, then snap-cooled on ice for 1 minute. The RNA was then mixed with 25 μL water and 10 μL 5X folding buffer (1.5 M sodium cacodylate pH 7.0, 50 mM MgCl_2_) and folded at 37 °C for 30 minutes. The folded RNA was modified by adding 5 μL of 15% dimethyl sulfate (v/v in ethanol) or water (negative control). Both reactions were incubated at 37 °C for 6 minutes, quenched by the addition of 50 μL beta-mercaptoethanol, purified using the Zymo RNA Clean and Concentrator 5 kit (Zymo Research), and eluted in 12 μL water. For reverse transcription, 10 μL of the modified RNA was mixed with 1 μL of 10 μM oVT555, incubated at 65 °C for 5 min, and snap cooled on ice. Then, the template-primer mix was combined with 4 μL of 5X TGIRT First Strand Synthesis Buffer (250 mM Tris-HCl pH 8.3, 375 mM KCl, 15 mM MgCl_2_), 2 μL 10 mM dNTPs, 1 μL freshly prepared 100 mM DTT, and 0.5 μL TGIRT-III (InGex LLC). The reaction was mixed and incubated at 57 °C for 3 hours. RNA was then hydrolyzed by addition of 10 μL hydrolysis buffer (0.5 M NaOH, 0.25 M EDTA) and incubation at 65 °C for 15 min. Hydrolysis was quenched by bringing the reaction volume to 50 μL with water, adding 100 μL of Oligo Binding Buffer (Zymo Research), and proceeding through the Zymo Oligo Clean and Concentrator (Zymo Research) purification protocol, eluting in 15 μL of water. 5 μL of the purified cDNA was amplified in a NEBNext Q5 HotStart master mix PCR reaction containing 0.5 μM each oVT554 and oVT555 with the following cycling conditions: 98 °C for 30 seconds, 10 cycles of 98 °C for 10 seconds followed by 72 °C for 60 seconds, with a final extension of 72 °C for 5 minutes. Products were purified using 0.9X Select-a-Size DNA Clean and Concentrator MagBeads (Zymo Research). Amplification of the full-length cDNA product was verified on an agarose gel stained with SYBRSafe (Invitrogen).

For SHAPE-based chemical mapping, 500 ng of RNA was brought to 12 μL in water and denatured at 95°C for 2 minutes followed by snap-cooling on ice for 2 minutes. The RNA was then folded by adding 6 μL of 3.3X SHAPE folding buffer (333 mM HEPES pH 8.0, 333 mM NaCl, 33 mM MgCl_2_) and incubating at 37°C for 20 minutes. 9 μL of the folded RNA was mixed with 1 μL of either 100 mM 1M7 (1-Methyl-7-nitroisatoic anhydride, freshly mixed in DMSO) or neat DMSO (negative control), mixed thoroughly by pipetting, and incubated at 37°C for 75 seconds (roughly 5 1M7 hydrolysis half-lives). After chemical treatment, the volume of both reactions was brought to 50 μL with water, cleaned up with the Zymo RNA Clean and Concentrator 5 kit, and eluted in 12 μL of water. 10 μL of eluted RNA was mixed with 1 μL of 200 ng/μL random nonamer primer (New England Biolabs), incubated at 65°C for 5 minutes, then snap-cooled on ice. Then, 8 μL of 2.5X MaP buffer (125 mM Tris pH 8.0, 187.5 mM KCl, 25 mM DTT, 1.25 mM each dNTPs, 15 mM MnCl2), was added to the primer-template mixture, incubated at room temperature for 2 minutes, and then mixed with 1 μL of SuperScript II (Invitrogen). The reverse transcription reaction was thoroughly mixed by pipetting, incubated at room temperature for 10 minutes, then at 42 °C for 3 hours. The reverse transcription enzyme was heat inactivated at 70°C for 15 min, snap-cooled on ice, then immediately mixed with 8 μL NEBNext Second Strand Synthesis Reaction Buffer (New England Biolabs), 4 μL NEBNext Second Strand Synthesis Enzyme Mix (New England Biolabs), and 48 μL water. The second strand synthesis reaction was mixed thoroughly through pipetting and incubated in a thermocycler with the heated lid off at 16°C for 60 min. The resulting double-stranded cDNA was purified with 1.8X volumes of Select-a-Size DNA Clean and Concentrator MagBeads (Zymo Research).

Double-stranded cDNA was prepared for Illumina sequencing with the NEBNext Ultra II FS DNA Library Prep Kit for Illumina (New England Biolabs) and iTru primers. Briefly, 100-500ng of DNA from either 1M7 or DMS conditions in 26 μL of water was fragmented and end-repaired by adding 7 μL NEBNext Ultra II FS Reaction Buffer and 2 μL NEBNext Ultra II FS Enzyme Mix, mixed thoroughly by vortexing, and incubated in a thermocycler with heated lid on at 37°C for 20 min, then 65°C for 20 min. Fresh ligation adapter was prepared by denaturing a solution of 15 μM each of iTrusR1-stub and iTrusR2-stubRCp in salty TLE (10 mM Tris pH 8.0, 0.1 mM EDTA, 100 mM NaCl) at 95°C for 1 minute, then annealed via slow cooling at −0.1°C/s to 25°C. The fragmented and end-repaired DNA was mixed with 2.5 μL 15 μM ligation adapter, 30 μL NEBNext Ultra II FS Ligation Master Mix, and 1 μL NEBNext Ultra II FS Ligation Enhancer. The ligation reaction was incubated in a thermocycler (heated lid off) at 20°C for 60 min. Ligated fragments were purified with 0.9X Select-a-Size DNA Clean and Concentrator MagBeads (Zymo Research) and eluted in 25 μL water. Dual-index sample barcodes were added through indexing PCR using the NEBNext Q5 Hot Start HiFi PCR Master Mix in a reaction with 0.5 μM each of iTru_5 and iTru_7 indexing primers with the following thermocycling parameters: 98°C for 30 s, 8 cycles of 98°C for 10 s, 55°C for 10 s, 72°C for 15 s, and a final extension at 72°C for 5 min. The final sequencing libraries were purified with 0.9X volumes of Select-a-Size DNA Clean and Concentrator MagBeads (Zymo Research), quantified by qPCR using the iTaq Universal SYBR Green Supermix (Bio-Rad) with iTru_P5 and iTru_P7 primers, pooled in equimolar concentrations, and sequenced on an Illumina Miseq (Stanford Protein and Nucleic Acid Core Facility) for 600 cycles.

Demultiplexed reads were downloaded from the BaseSpace Sequence Hub (Illumina). Most analysis steps were carried out using the RNAFramework RNA structure probing analysis toolkit^92^. Using the ‘rf-map’ module, reads were trimmed with CutAdapt^88^ and mapped to the wild-type RNA sequence using Bowtie 2^89^. Mutations were counted using the ‘rf-count’ module with ‘-m’ flag, then normalized using the ‘rf-norm’ module with ‘-sm 4 -nm 2 -rb AC -nw 50 -dw’ flags for DMS samples and ‘-sm 3 -nm 2 -rb N -nw 50 -dw’ flags for 1M7 samples. These normalized reactivities (**Table S4**) were plotted as-is, and also used to predict RNA secondary structure using the RNAStructure^93^ ‘Fold’ command, implemented in Arnie (https://github.com/DasLab/arnie).

### High-throughput in-line and SHAPE probing on Eterna-designed RNA fragments (In-line-seq)

The In-line-seq experiments relied on a different pipeline for massively parallel RNA generation, treatment, and Illumina sequencing than mRNA experiments above. We describe these steps below; see ref.^94^ for further details and **Table S6** for primers.

#### Preparation of DNA templates

DNA fragments encoding for RNA molecules from the Eterna ‘OpenVaccine: Roll-your-own-structure’ challenge were ordered in the form of a custom oligonucleotide pool of DNA (Custom Array/Genscript) with the 20-nt T7 RNA polymerase promoter sequence (5’-TTCTAATACGACTCACTATA-3’) prepended to each DNA. Amplification of the DNA template was performed via emulsion PCR. A hydrophobic solution (‘oil phase’) containing 80 μL of ABIL EM90 (Evonik Corporation), 1 μL of Triton X-100, and 1919 μL of mineral oil was vortexed for 5 minutes and then incubated on ice for 30 minutes. Then, 75 μL of water-soluble reaction mixture (liquid phase) containing 1X Phire Hot Start II buffer, 0.2 mM dNTPs, 1.5 μL of Phire II DNA polymerase, 2 μM of each primer (T7 promoter and Tail2 Reverse complement), 0.5 mg/ml of BSA, and 360 ng of the oligonucleotide pool was prepared. In a 1.0 ml glass vial (kept on ice and frozen at −20°C overnight before use), 300 μL of the oil phase was added into the glass vial and vortexed at 1000 rpm for 5 minutes. Next, 10 μL of the liquid phase was added followed by 10 seconds of vortexing. This addition of the liquid phase followed by vortexing was repeated 4 times such that 50 μL of the liquid phase has been added to 300 μL of the oil phase in the vial. The now-mixed 350 μL of emulsion PCR solution was the subjected to the following thermocycling protocol: 98°C for 30 seconds for initial denaturation, 42 cycles of amplification (98°C for 10 seconds, 55°C for 10 seconds, and 72°C for 30 seconds), and final extension at 72°C for 5 minutes

The PCR was purified by adding 100 μL of mineral oil, followed by a brief vortex (~10 seconds) and centrifugation at 13,000 rcf for 10 minutes. The oil phase was then discarded. 1 mL of diethyl ether was added, followed by a brief vortex (~10 seconds) and centrifugation at 13,000 rcf for 1 minute. The upper layer (termed the detergent layer) was then discarded. Extraction with diethyl ether was repeated 3 times to ensure through purification. The resulting product was then incubated at 37°C for 5 minutes, adjusted to a final volume of 40 μL with nuclease-free water, then purified with 72 μL of Ampure bead XP (Beckman Coulter) following the standard protocol specified by the vendor. Finally, DNA was eluted into 20 μL of nuclease-free water.

#### Preparation of RNA templates

A library of RNA molecules was then prepared from the amplified DNA template using the TranscriptAid T7 High Yield Transcription Kit (Thermofisher, K0441) using the reaction mixture specified by the vendor. Transcription was performed with incubation at 37°C for 3 hours. After transcription, the DNA template was removed through the addition of 2 μL of DNAse I (add vendor) followed by incubation at 37°C for 30 minutes. After DNA digestion, the RNA was purified using a mixture of AMPure XP beads (Beckman Coulter) with 40% polyethylene glycol (mixed in a 7:3 ratio). Final elution into 25 μL of nuclease-free water yielded purified RNA.

#### Degradation in-line probing of RNA samples

For degradation experiments, 45-50 pmol of RNA was subject to four conditions: 1) 50 mM Na-CHES buffer (pH 10.0) at room temperature without added MgCl_2_; 2) 50 mM Na-CHES buffer (pH 10.0) at room temperature with 10 mM MgCl_2_; 3) phosphate buffered saline (PBS, pH 7.2; Thermo Fisher Scientific-Gibco 20012027) at 50°C without added MgCl_2_; and 4) PBS (pH 7.2) at 50°C with 10 mM MgCl_2_.

For degradation reactions containing MgCl_2_, RNA was collected at 0 and 24 hour time points. For reactions without MgCl_2_, timepoints were collected at 0 and 7 days. At each timepoint, the degradation reaction was quenched with 15 μL of 500 mM of Tris-HCl (pH 7) and 3 μL of 500 mM of Na-EDTA. Quenched samples were brought to a final volume of 100 μL with nuclease-free water. The RNA was purified through precipitation as follows. 1.5 μL of Glyco Blue (Thermo Fisher), 10 μL of 3 M sodium acetate (pH 5.2), and 330 μL of cold 100 % ethanol were added, the reaction mixture mixed, and incubated on dry ice for 20 to 30 minutes. After incubation, the reaction was centrifuged at 21,000 rcf for 30 minutes. The resulting pellet was washed twice with 500 μL of cold 70% ethanol, and pelleted after each wash step. Finally, the reaction was dried for 10 minute at room temperature to remove any residual ethanol, and resuspended in 5 μL of nuclease-free water.

#### Structure probing of RNA samples

In parallel with the in-line hydrolytic degradation conditions above, we carried out SHAPE structure probing experiments^64^, as follows. 15 pmol of purified RNA was added to 2 μL of 500 mM HEPES buffer (pH 8.0) and denatured at 90°C for 3 minutes. The reaction was then cooled down to room temperature over 10 minutes. 2 μL of 100 mM MgCl_2_ was then added, followed by incubation at 50°C for 30 minutes. The sample was cooled down to room temperature over 20 minutes before addition of 5 μL of 1-methyl-7-nitroisatoic anhydride (1M7, 8.48 mg/mL of DMSO) followed by incubation at room temperature for 15 min, and brought to a final volume of 20 μL with nuclease-free water. The reaction was quenched with 5 μL of 500 mM Na-MES pH 6.0, the reaction was adjusted to be 100 μL, and purified with ethanol precipitation as above. As a control, a sample was prepared in parallel without addition of 1M7 but subject to the same protocol described above.

#### Preparation of cDNA and sequencing

cDNA was prepared from the six RNA samples (two from structure probing, four from degradation). 5 μL of purified RNA was added to a reaction mixture containing 1x First Strand buffer (Thermo Fisher), 5 mM dithiothreitol (DTT), 0.8 mM dNTPs, 0.6 μL of SuperScript III RTase (Thermo Fisher) to a final volume of 15 μL. The reaction was incubated at 48 °C for 40 minutes and stopped with 5 μL of 0.4 M sodium hydroxide. The reaction was then incubated at 90 C for 3 minutes, cooled on ice for 3 minutes, and neutralized with 2 μL of quench mix (prepared as 2 mL of 5 M sodium chloride, 3 mL of 3 M sodium acetate, 2 mL of 2 M hydrochloric acid).

cDNA was pooled down with 1.5 uL of Oligo C’ beads^64^ (in house magnetic beads prepared by immobilizing 2x Biotin oligonucleotides with Dynabeads™ MyOne™ Streptavidin C1; Thermo Fisher Scientific 65001), washed twice with 70% ethanol, then resuspended in 3.0 μL of water. We pooled 1.5 μL of each sample together, and took 9 μL of cDNA to continue to ligation an Illumina adapter by using Circ. Ligase I (Lucigen) at 68°C for 2 h. The reaction was stopped by incubation at 80 °C for 10 min. cDNA was added to 10 ul of 5 M NaCl and pulled down with a magnetic stand and washed with 70% ethanol; the ligated product was resuspended in 15 μL H_2_O.

Ligated product was quantified by qPCR. dsDNA at 3 nM concentration was sequenced using an Illumina Miseq (High output, Read 1 = 101 cycles and Read 2 = 51 cycles). The resulting data were analyzed using MAPseeker (https://ribokit.github.io/MAPseeker)^95^ following the recommended steps for sequence assignment, peak fitting, background subtraction of the no-modification control, correction for signal attenuation, and reactivity profile normalization.

### In-line and SHAPE probing by capillary electrophoresis

One-by-one followup to profile degradation of RNA fragments at single-nucleotide resolution through capillary electrophoresis was carried out with some differences to the pooled In-line-seq experiments above. We describe these steps below; see ref.^94^ for further details and **Table S6** for primers.

#### DNA template preparation

DNA templates were designed to include the 20-nt T7 RNA polymerase promoter sequence (5’-TTCTAATACGACTCACTATA-3’) followed by the remaining sequence encoding desired RNA. Double-stranded templates were prepared by extension of 60-nt DNA oligomers (IDT, Integrated DNA Technologies) with Phusion DNA polymerase (Finnzymes), using the following thermocycler protocol: denaturation for 30 sec at 98°C, 35 cycles of denaturation for 10 sec at 98°C annealing for 30 sec at 60 to 64°C , extension for 30 sec at 72°C; final extension for 10 min at 72 °C and cooling to 4°C.

DNA samples were purified with AMPure magnetic beads (Agencourt, Beckman Coulter) following manufacturer’s instructions. Sample concentrations were estimated based on UV absorbance at 260 nm measured on Nanodrop 100 or 8000 spectrophotometers. Verification of template length was accomplished by electrophoresis of all samples and 10-bp and 20-bp ladder length standards (Thermo Scientific O’RangeRuler SM1313 & SM1323) in 4% agarose gels (containing 0.5 mg/mL ethidium bromide) and 1x TBE (100 mM Tris, 83 mM boric acid, 1 mM disodium EDTA). All sample manipulations, including following steps, were carried out in 96-well V-shaped polypropylene microplates (Greiner).

#### Preparation of RNA templates

*In vitro* transcription reactions were carried out in 40 μL volumes with 10 pmol of DNA template, using the TranscriptAid T7 High Yield Transcription Kit (Thermo Fisher, K0441). In some syntheses, pseudouridine-5’-triphosphate (TriLink Biotechnologies, N-1019) or N1-methylpseudouridine-5’-triphosphate (TriLink Biotechnologies, N-1081) were used to replace regular UTP. Reactions were incubated for 3 hours at 37°C, followed by degradation of DNA template with 2 μL of DNase I at 37°C for 30 min. RNA samples were purified with 1.8x volume of AMPure XP beads (Beckman Coulter) mixed with 40% PEG-8000 (ratio of 7:3), following manufacturer’s instructions. Concentrations were measured by absorbance at 260 nm on Nanodrop 100 or 8000 spectrophotometers.

#### In-line probing (hydrolytic degradation profiling)

For degradation experiments, 750 ng of RNA was subjected to 50 mM Na-CHES buffer (pH 10) at room temperature with 10 mM MgCl_2_ at a final volume of 30 μL. RNA was collected at 0- and 24-hour time points. At each timepoint, 15 μL of reaction was taken, and degradation quenched with 7.5 μL of 500 mM of Tris-HCl (pH 7) and 1.5 μL of 500 mM of Na-EDTA. Quenched samples were purified with 2 μL of Oligo dT bead (Thermo Fisher, AM1922) and 0.8 μL of 10 μM FAM-A20-Tail2. RNA, bead, and primer were incubated at room temperature for 15 min, pulled down by magnetic stand for 10 min, and washed twice with 70% ethanol and left to dry on the bead. RNA was eluted in 2.5 μL of nuclease-free water (RNA was binded with OligodT bead and FAM.A20 Tail2 at this point). Subsequently, these RNAs were further treated and purified with steps analogous to SHAPE probing (see next) to enable side-by-side comparisons. RNAs were added to 2 μL of 500 mM Na-HEPES buffer (pH 8.0) and denatured at 90°C for 3 minutes. The reaction was then cooled down to room temperature over 10 minutes. 2 μL of 100 mM MgCl_2_ was then added, followed by incubation at 50°C for 30 minutes. The sample was cooled down to room temperature over 20 minutes before addition of 5 μL of nuclease-free water, followed by incubation at room temperature for 15 min, and brought to a final volume of 20 μL with nuclease-free water. The RNA sample was further purified by incubating the sample with 5.0 μL of Na-MES, pH 6.0, 3.0 μL of 5 M NaCl, and brought to a final volume of 10 μL with nuclease-free water. The reaction mixture was incubated at room temp for 15 min, pulled down by 96-post magnetic stand for 10 min, washed twice with 70% ethanol and allowed to dry, before adding 2.5 μL of nuclease-free water.

#### SHAPE experiments

For SHAPE structure probing experiments, 1.2 pmol of purified RNA was added to 2 μL of 500 mM Na-HEPES buffer (pH 8.0) and denatured at 90°C for 3 minutes. The reaction was then cooled down to room temperature over 10 minutes. 2 μL of 100 mM MgCl_2_ was then added, followed by incubation at 50°C for 30 minutes. The sample was cooled down to room temperature over 20 minutes before addition of 5 μL of nuclease-free water (negative control) or 1-methyl-7-nitroisatoic anhydride (1M7, 8.48 mg/mL of DMSO) followed by incubation at room temperature for 15 min, and brought to a final volume of 20 μL with nuclease-free water. The SHAPE-RNA sample was further purified by incubating the sample with 5.0 μL of Na-MES, pH 6.0, 3.0 μL of 5 M NaCl, 1.5 μL of Oligo dT bead, 0.25 μL of 10 μM FAM-A20-Tail2, and brought to a final volume of 10 μL with nuclease-free water. The reaction mixture was incubated at room temp for 15 min, pulled down by 96-post magnetic stand for 10 min, washed twice with 70% ethanol and allowed to dry, before adding 2.5 μL of nuclease-free water.

#### Preparation of in-line probing and SHAPE samples for capillary electrophoresis

cDNA was prepared from in-line probing and SHAPE RNA samples as follows (note that above procedures leave RNA bound to FAM-A20-Tail2 reverse transcription primers which are in turn bound to Oligo dT beads). 2.5 μL of purified RNA was added to a reaction mixture containing 1x First Strand buffer (Thermo Fisher), 5 mM dithiothreitol (DTT), 0.8 mM dNTPs, 0.2 μL of SS-III RTase (Thermo Fisher) to a final volume of 5.0 μL. The reaction was incubated at 48°C for 40 minutes, and stopped with 5 μL of 0.4 M sodium hydroxide. The reaction was then incubated at 90°C for 3 minutes, cooled on ice for 3 minutes, and neutralized with 2 μL of quench mix (2 mL of 5 M sodium chloride, 3 mL of 3 M sodium acetate, 2 mL of 2 M hydrochloric acid). For four cDNA reference ladders, each of four ddNTPs (GE Healthcare 27-2045-01) with a ddNTP/dNTP ratio of 1.25 (0.1 mM / 0.08 mM) was used in the reverse-transcription reaction.

cDNA was pulled down on a 96-post magnetic stand and washed 2 times with 100 μL 70% ethanol. To elute the bound cDNA, the magnetic beads were resuspended in 10.0625 μL ROX350 (Thermo Fisher Scientific 401735) /Hi-Di (0.0625 μL of ROX 350 ladder in 10 μL of Hi-Di formamide) and incubated at room temperature for 20 minutes. The cDNA was further diluted by 1/3 and 1/10 in ROX350/HiDi and samples loaded onto capillary electrophoresis sequencers (ABI-3730) on capillary electrophoresis (CE) services rendered by ELIM Biopharmaceuticals. CE data was analyzed using the HiTRACE 2.0 package^96^ (https://github.com/ribokit/HiTRACE), following the recommended steps for sequence assignment, peak fitting, background subtraction of the no-modification control, correction for signal attenuation, and reactivity profile normalization.

### Measurement of in-solution mRNA stability by capillary electrophoresis

For one-by-one measurement of in-solution mRNA stability, *in vitro* transcribed mRNA was incubated in a degradation buffer over ten time points (0, 0.5, 1.0, 1.5, 2, 3, 4, 5, 18, and 24 hours), then analyzed by capillary electrophoresis.

For each time point, 1.6 pmol of mRNA brought to 10 μL in a buffer containing 50 mM Na-CHES at pH 10 with 10 mM MgCl_2_, and the reaction was incubated at 25 °C. When the incubation period was reached for each time point, 5 μL of Tris-HCl at pH 7 and 1 μL of 500 mM EDTA in nuclease free water was added to quench the degradation reaction, and frozen for further analysis. After the final time point (24 hours), 4 μL of each mRNA degradation sample (out of a total stored volume of 16 μL) was taken, and mixed with 1 μL of a control RNA at a concentration of 50 ng/μL. For these experiments we opted to use the P4-P6 domain of the *Tetrahymena* ribozyme with two added hairpins^94^ (~239 nt) as our control. The RNA mixture was then purified using a mixture of AMPure XP beads (Beckman Coulter) with 40% polyethylene glycol (mixed in a 7:3 ratio). The resulting RNA was eluted into 4.5 μL of RNAse-free water for analysis on the 2100 Bioanalyzer (Agilent) using the RNA-Nano Eukaryote protocol.

The data from the Bioanalyzer were analyzed using a custom script that performs the following analysis. We first converted elution times to nucleotides based on a ladder control (25, 200, 500, 1000, 2000, and 4000 nts). We then estimated relative mRNA amounts based on peak areas at expected band lengths (in our case, ~900 nucleotides for the mRNAs of interest and ~265 nucleotides for the control). When calculating peak areas, we performed background subtraction, where the background was defined as the area under a linear line in the range of nucleotides used for the peak area. Normalization was performed using two different methods used to cross-validate. First, we normalized the peak areas of full-length mRNA to the control P4-P6 domain RNA that was spiked into the samples after degradation was performed. Second, we also normalized peak areas of full-length mRNAs to the total amount of RNA in the lane less the peak area of the bands of interest (between ~20-1000 nucleotides in our case), assuming that the majority of the other RNA in the lane were degradation products from the mRNA of interest. These distinct approaches to normalizing the data gave the same results within estimated error (see below). After calculations of normalized peak areas, we then calculated fraction intact values for each mRNA by dividing the normalized area across the ten timepoints by the normalized area at the start (0 hours).

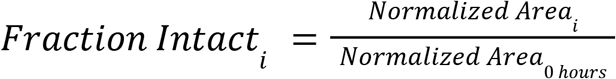

For each sample, we fit fraction intact values across the different timepoints to an exponential function,

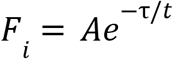

Where *F_i_* is an array of fraction intact values across multiple time points, *A* is the amplitude of the exponential decay function, τ is the time constant, and *t* is an array of time points in hours. We then used the time constant to calculate the *in vitro* half-life of mRNA:

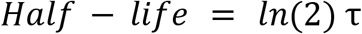

### Polysome selection and library preparation

The variant 5’ UTR is composed of: fixed first 29 nt of *hHBB*, variable 35 nt (initially degenerate) and 6 nt Kozak consensus. See **Table S6** for the detailed construct layout. To generate the reporter mRNA pool containing the variant 5’ UTR library, IVT template was first assembled by PCR under the following conditions: 4 μL 10x AccuPrime Pfx Reaction Mix, 0.4 pmol HBB29_N35 amplicon, 0.4 pmol Nluc_HBB_3UTR, 0.4 μL AccuPrime Pfx Polymerase in 40 μL of total reaction volume. Cycling conditions are: 95°C for 120 sec, and 19 cycles of 95°C for 15 sec, 66°C for 30 sec, 68°C for 75 sec. PCR product was purified on silica columns (NEB T1034) and amplified with under the following conditions: 4 uL 10X AccuPrime Pfx Reaction Mix, 4 μL 10 μM T7_28_HBB_30_F, 4 μL 10 μM Nanoluc_ORF_R, 0.4 μL AccuPrime Pfx Polymerase in 40 μL total reaction volume. Cycling conditions are: 95°C for 120 sec, and 4 cycles of 95°C for 15 sec, 66°C for 30 sec, 68°C for 75 sec. The mRNA was *in vitro* transcribed, capped and polyadenylated as described above.

Transfection of HEK-293 cells and sucrose gradient fractionation were performed as described above. Equal volumes of fractions 10-16 were pooled and RNA was by acidic phenol chloroform extraction followed by column purification (Zymo Research, R1013) as described above. 1/3 lysate volume was kept as input before layering onto the sucrose gradient and RNA was extracted from the input lysate by Trizol extraction followed by column purification. 1.5 μg RNA in 5.5 μL was mixed with 0.5 μL 2uM RT_Nluc26_UMI12_Read1Partial (**Table S6**) and 0.5 μL 10 mM dNTPs each. The RNA samples were then denatured at 65°C for 5 min and chilled to 4°C. 3.5 μL reverse transcription mix was added to 10 μL total reaction volume: 2 μL 5x Superscript IV buffer, 0.5 μL 10 mM DTT, 0.5 μL Superase-In (ThermoFisher Scientific, AM2694), 0.5 μL Superscript IV (Thermo 18091050). The reaction was incubated at 55°C for 45 min and inactivated at 80°C for 10 min. Variant 5’ UTR amplicon was amplified from the reverse transcription reaction via PCR under the following reaction conditions: 4 μL RT reaction, 40 μL 2x Q5 Hot Start Master Mix (NEB M0494S), 0.8 μL 100x SYBR (Thermo S7563), 4 μL 10 μM T7_28_HBB_29_F, 4 μL 10 μM Nanoluc_ORF_R, in 80 μL total reaction volume. Cycling conditions were as follows: 98°C for 60 sec, and 15 cycles of 98°C for 10 sec, 68°C for 10 sec, 72°C for 10 sec. PCR product was purified on silica columns (NEB T1034) and assembly with Nluc_HBB_3UTR fragment was performed as described above for initial preparation of IVT template using HBB29_N35 amplicon. The mRNA was *in vitro* transcribed, capped and polyadenylated as described above. The same process of transfection, fractionation, reverse transcription, PCR amplification, assembly and in vitro transcription was repeated.

For sequencing library preparation the RT reaction was PCR amplified under the following conditions: 1 μL RT reaction, 10 μL 2x Q5 Hot Start Master Mix (NEB M0494S), 0.2 μL 100x SYBR (Thermo S7563), 1 μL 10 μM Read1, 1 μL 10 μM Read2Partial_HBB29 in 20 μL total reaction volume. Cycling conditions were as follows: 98°C for 60 sec, and 15 cycles of 98°C for 10 sec, 68°C for 10 sec, 72°C for 10 sec. Sequencing adaptors were added using the following conditions for final round PCR: 1 μL first round PCR reaction, 10 μL 2x Q5 Hot Start Master Mix, 0.2 μL 100x SYBR, 1 μL 10uM NEBNext Index Primer (NEB E7335, NEB E7500, NEB E7710, NEB E7730, NEB E6609), 1μL 10uM NEBNext Universal PCR Primer in 20 μL total volume. Cycling conditions are: 98°C for 60 sec, and 5 cycles of 98°C for 10 sec, 72°C for 10 sec. All barcoded samples were then pooled at equal volumes and purified with 1.1x SPRIselect beads Beckman Coulter B23317). Sequencing was performed at the Stanford Functional Genomics Facility (SFGF) at Stanford University, on the Illumina NextSeq 550 instrument, using a high output kit, 1×81 cycles. Primer sequences and the sequencing construct layout are provided in **Table S6**.

### Polysome selection library sequencing data analysis

Following adapter trimming, 670440 sequences with at least 10 summed read count across all libraries combined were set as the reference. Each library was aligned to this indexed reference using Bowtie2^89^. Only uniquely mapping reads with edit distance ≤3 were retained. Alignments were further deduplicated using UMIcollapse (-p 0.05, -k 1)^90^. This results in the matrix of read count where rows are different sequence variants and columns are the samples.

Normalized counts were obtained by dividing the matrix column-wise by total read counts per sample. For sequence variants with at least 15 reads in any one of the samples, a regression model was fitted on normalized read counts with the sequential selection rounds as ordinal predictors, penalizing differences between coefficients of adjacent groups (R package ordPens) ^97^. False discovery rate was estimated by Benjamini-Hochberg procedure. For choosing the final set of candidates, the criteria of ≥15 read counts in the final round polysome selection library and ≥2 fold enrichment over input in the final round was also required.

For analysis of k-mers, 1 million reads (prior to alignment and analysis for highly enriched individual sequences) were sampled from each library. Position-specific k-mers were counted and statistical significance of pair-wise enrichments for each position-specific k-mers are calculated using kpLogo^85^. The parameters were: zero-order Markov model background; 2≤k≤6; -shift 0 and -max-shift 0; binomial test with Bonferroni correction.

### Design algorithms/analysis

The protein sequences for each target were used to generate DNA sequences at Integrated DNA Technologies (IDT, https://www.idtdna.com/CodonOpt), Twist Biosciences (https://ecommerce.twistdna.com/app), and GENEWIZ (https://clims4.genewiz.com/Toolbox/CodonOptimization). Solutions from LinearDesign were obtained using the LinearDesign server (http://rna.baidu.com/). Ribotree solutions were terminated after 6000 iterations. The RiboTree code is available for noncommercial use at https://eternagame.org/about/software, with example usage to replicate runs for this work provided.

Structure prediction and ensemble-based calculations were performed using LinearFold and LinearPartition with ViennaRNA, CONTRAfold, and EternaFold parameters. Secondary structure features were calculated from predicted MFE structures using RiboGraphViz (www.github.com/DasLab/RiboGraphViz). CAI (Codon Adaptation Index) was calculated as the geometric mean of the relative usage frequency of codons along the length of the coding region, as previously described^45^:

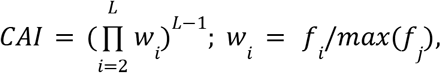

Where f_j_ represents the frequency of all codons coding for amino acid at position i. Scripts to reproduce degradation and structure predictions are available in the “OpenVaccine-solves” database under an Open COVID license at https://eternagame.org/about/software.

### Quantification and Statistical Procedures

In all figures, data are presented as mean, SD or SEM as stated in the figure legends, and *p ≤ 0.05 is considered significant (ns: p > 0.05; *p ≤ 0.05; **p ≤ 0.01; ***p ≤ 0.001; ****p ≤ 0.0001). Blinding and randomization were not used in any of the experiments. Number of independent biological replicates used for the experiments are listed in the figure legends. Tests, two-tailed unpaired Student’s t-test if not stated otherwise, and specific p-values used are indicated in the figure legends. In all cases, multiple independent experiments were performed on different days to verify the reproducibility of experimental findings. Errors for in-cell degradation coefficients are standard errors of coefficient estimates. Errors for in-cell ribosome load are estimated by bootstrapping where fraction labels are shuffled and before scaling by spike-in normalization factors. Errors for predicted expression values are estimated by Taylor series method. Error and significance tests for fitting single-exponential in-solution half-lives were estimated by non-parametric bootstrapping.

### ACCESSION NUMBERS

RNA sequencing data from RNA-seq experiments are being deposited in the Gene Expression Omnibus (GEO). Single-nucleotide-resolution in-line probing and SHAPE data are deposited at the following RNA Mapping Database^98^ (http://rmdb.stanford.edu) accession numbers:

In-line-seq datasets:

RYOS1_NMD_0000 (no modification)
RYOS1_MGPH_0000 ([Mg2+] = 10 mM, pH = 10, 24°C, 1 day)
RYOS1_PH10_0000 ([Mg2+] = 0 mM, pH = 10, 24°C, 7 days)
RYOS1_MG50_0000 ([Mg2+] = 10 mM, pH = 7.2, 50°C, 1 day)
RYOS1_50C_0000 ([Mg2+] = 0 mM, pH = 7.2, 50°C, 7 days)
SHAPE_RYOS_0620 (SHAPE 1M7 reactivity)

One-by-one follow ups

RYOSFL_MOD_0001
RYOSFL_MOD_0002
RYOSFL_MOD_0003
RYOSFL_MOD_0004
RYOSFL_MOD_0005
RYOSFL_MOD_0006
RYOSFL_MOD_0007
RYOSFL_MOD_0008

## SUPPLEMENTAL INFORMATION

Supplemental Information includes 7 Supplemental Figures and 7 Supplemental Tables.

## SUPPLEMENTAL TABLES

**Table S1.** Attributes for pooled 233 sequences, including nucleotide sequences, sequence annotation, ribosome load, in-solution RNA degradation half-lives, in-cell RNA degradation half-lives, predicted protein expression, and biophysical properties. “NA” indicates values not measured or calculated for select constructs.

**Table S2.** Attributes for CoV-2 5’ UTR mutagenesis sequences

**Table S3.** Frequency and statistics of position-specific k-mers after polysome selection.

**Table S4.** Normalized DMS and SHAPE (1M7) data on P4-P6, Yellowstone and LinearDesign-1 sequences.

**Table S5.** Attributes for 24 CDS designs, including designer/source, nucleotide sequences, in-solution half-lives, luciferase expression values, biophysical properties, and predicted structural metrics.

**Table S6.** List of primers and construct layouts.

**Table S7**. Eterna OpenVaccine participants List of primers and construct layouts

**Table S8**. List of OpenVaccine donors.

